# Glutamate indicators with improved activation kinetics and localization for imaging synaptic transmission

**DOI:** 10.1101/2022.02.13.480251

**Authors:** Abhi Aggarwal, Rui Liu, Yang Chen, Amelia J Ralowicz, Samuel J Bergerson, Filip Tomaska, Timothy L Hanson, Jeremy P Hasseman, Daniel Reep, Getahun Tsegaye, Pantong Yao, Xiang Ji, Marinus Kloos, Deepika Walpita, Ronak Patel, Manuel A Mohr, Paul W Tilberg, Boaz Mohar, The GENIE Project Team, Loren L Looger, Jonathan S Marvin, Michael B Hoppa, Arthur Konnerth, David Kleinfeld, Eric R Schreiter, Kaspar Podgorski

**Author notes:** Correspondence should be addressed to K.P.

## Abstract

The fluorescent glutamate indicator iGluSnFR enables imaging of neurotransmission with genetic and molecular specificity. However, existing iGluSnFR variants exhibit saturating activation kinetics and are excluded from post-synaptic densities, limiting their ability to distinguish synaptic from extrasynaptic glutamate. Using a multi-assay screen in bacteria, soluble protein, and cultured neurons, we generated novel variants with improved kinetics and signal-to-noise ratios. We also developed surface display constructs that improve iGluSnFR’s nanoscopic localization to post-synapses. The resulting indicator, iGluSnFR3, exhibits rapid non-saturating activation kinetics and reports synaptic glutamate release with improved linearity and increased specificity versus extrasynaptic signals in cultured neurons. In mouse visual cortex, imaging of iGluSnFR3 at individual boutons reported single electrophysiologically-observed action potentials with high specificity versus non-synaptic transients. In vibrissal sensory cortex Layer 4, we used iGluSnFR3 to characterize distinct patterns of touch-evoked feedforward input from thalamocortical boutons and both feedforward and recurrent input onto L4 cortical neuron dendritic spines.

## Introduction

Glutamate is the most abundant neurotransmitter molecule in the vertebrate brain, with approximately one glutamatergic synapse per cubic micrometer of neuropil (1, 2). Despite such close packing, synaptic glutamate signaling incurs little crosstalk (3). Synapses achieve this remarkable spatial specificity partly through the kinetics and localization of ionotropic glutamate receptors. Glutamate is released when synaptic vesicles fuse to the presynaptic membrane, producing brief and highly localized concentration transients that are sensed by closely-apposed post-synaptic glutamate receptors with low affinity and rapid activation kinetics (4, 5). Synaptic glutamate is quickly taken up by glutamate transporters, but some escapes the synaptic cleft and signals extra-synaptically via high-affinity receptors (3). Methods are needed to distinguish synaptic from extrasynaptic glutamate signals, and to monitor glutamate with the same high spatial specificity achieved by synaptic glutamate receptors. For example, the ability to specifically detect inputs from a neurons’ synaptic partners is needed to understand the nature of dendritic integration, including the spatial arrangement of synaptic inputs (6, 7), mechanisms that govern synaptic plasticity(8, 9), and the transformations that neurons perform on their inputs (10, 11). Linear and rapid measurements are needed to monitor the amount and timing of glutamate release, and high signal-to-noise ratios are needed to monitor release under challenging conditions, such as in deep structures in vivo over single behavioral trials.

The intensity-based Glutamate-Sensing Fluorescent Reporter (iGluSnFR) and its variants have been widely adopted for neural glutamate measurements. iGluSnFR has been used to image synaptic release, including spontaneous and evoked quantal release (12–16), as well as extra-synaptic signaling and glutamate clearance (17, 18). However, the ability of commonly-used iGluSnFR variants to distinguish synaptic from extra-synaptic signals has not been established. Experiments have measured or bounded the spatial extent of iGluSnFR and SF-iGluSnFR signals under multiple conditions (10,13,18–21), but these bounds have been larger than the mean inter-synaptic distance. For example, SF-Venus-iGluSnFR.A184S exhibited signal correlations spanning 3.6 micrometers in visual cortex *in vivo* (18), corresponding to a volume containing roughly 30 glutamatergic synapses (1, 2). It has therefore been unclear, on the basis of iGluSnFR measurements alone, whether a given signal originates from a labeled neuron’s own synapse or from nearby unconnected axons. Based on measurements of indicator ON rates (20, 22), we realized that the specificity of common iGluSnFR variants for synaptic versus extra-synaptic signals is limited by *K_M_*, the rate-limiting glutamate concentration. Existing variants have low values of *K_M_*, meaning that the indicator ON rate is saturated during high-concentration glutamate transients and responses are therefore reduced at release sites (“kinetic saturation”; Supplemental Note 1). Experiments also suggested that the existing approach for displaying iGluSnFR on cell surfaces using the PDGFR transmembrane domain results in poor nanoscopic localization at post-synapses; iGluSnFR signals have generally been larger at axonal boutons than on dendrites (20) and fusion to a post-synaptically-localized protein dramatically increased signals of a low-affinity variant (13).

Here we set out to address these issues and engineer new sensitive iGluSnFR variants with improved specificity for synaptic versus extrasynaptic glutamate. We performed a screen involving 20 generations of diversification and selection in bacteria (∼10^6^ variants screened) and lysate (∼10^4^ variants screened), with the final two generations including assays in cultured neurons (∼10^3^ variants screened), while selecting for kinetic and photophysical properties predicted to improve synapse-specific signals. With the winning variant, iGluSnFR3.v857, we performed a second screen of C-terminal membrane anchoring domains to identify variants with improved synaptic responses in neuronal cell culture, then investigated their nanoscopic localization to post-synapses in vivo. We assessed the linearity and synaptic specificity of iGluSnFR3.v857 in neuronal cell culture by manipulating vesicle release. In the mouse cortex in vivo, we characterized iGluSnFR variants’ responses to single presynaptic action potentials using simultaneous patch-clamp electrophysiology and imaging, and responses to controlled sensory stimulation across temporal frequencies. Finally, we used iGluSnFR3.v857 to compare the spatiotemporal activity patterns of feedforward thalamocortical and recurrent inputs to layer 4 (L4), 400 micrometers below the pia. We showed that thalamocortical boutons exhibit low-latency responses phase-locked to whisker stimulation, while spines of L4 excitatory cortical neurons in the same compartment each exhibit characteristic timings that vary over tens of milliseconds, reflecting the timing distribution of recurrent inputs. The measurement of these distinct patterns on densely intertwined neuronal structures demonstrates the high spatial specificity of iGluSnFR3.v857 for synaptic versus extrasynaptic glutamate.

## Results

### Generation of iGluSnFR3 variants through multi-assay screening

As the template for our screen we picked SF-Venus-iGluSnFR-A184V (hereafter, WT), a yellow variant with the highest K_M_ among superfolder variants, and a larger two-photon (2P) cross-section than green iGluSnFR variants (20), especially at excitation wavelengths longer than 1 micrometer better for co-excitation with red fluorophores and deep in vivo imaging (23). We evolved a population of indicators through 20 rounds of mutagenesis and screening in bacteria, purified protein, and cultured neurons (Supplemental Note 2). We used high-throughput assays to rapidly discard unwanted variants, followed by lower-throughput assays on remaining variants, to richly characterize variants in each round without compromising library size.

In each round, the input population was first diversified by either random mutagenesis, recombination, or site-directed mutagenesis to create a library of soluble iGluSnFR variants. We first screened bacterial colonies (∼10^5^ per round) under 2-photon (rounds 5-8) or 1-photon (other rounds) excitation, selecting the brightest colonies to retain variants that express and mature efficiently and have high glutamate-bound brightness ([Glu]_cytosolic_≈100 mM in *E*.

*coli*). We assayed these variants (∼400 per round) in clarified bacterial lysate, where we measured fluorescence spectra before and after adding glutamate (200-500 μM, depending on round), retaining only those with large responses and desirable spectral properties. For the 4-20 best variants per round, we measured (in various rounds): affinity, pH response, quantum yield, extinction coefficient, 2-photon spectra, fluorescence correlation spectra, DNA sequence, and kinetic parameters (see Methods). In the final 2 rounds, a larger number of variants were selected, cloned into a surface display vector (pMinDisplay), then expressed and imaged in cultured rat neurons (24). We measured brightness, membrane trafficking, bleaching, and responses to 1 and 20 field-stimulated action potentials (APs) using high-speed (180 frames/s) imaging to assess kinetics and spatial specificity (Supplemental Note 2). Selected variants were then diversified and returned to the bacterial screen before the next round of neuronal screening. From this process we selected the variant iGluSnFR3.v857 (“v857”, the 857th variant screened in neurons; 15 mutations vs WT; Supplemental Note 3) on the basis of its high K_M_, rapid on-cell kinetics, high brightness, large fractional fluorescence response (ΔF/F_0_), and high photostability (Figure 1a-e, Figure 2a-e). We selected the closely-related variant iGluSnFR3.v82 (“v82”, the 82nd variant screened in neurons; 13 mutations vs WT) for comparison, due to its large 1 AP response amplitude and higher affinity, but kinetic properties less favorable for synaptic specificity than v857.

**Figure 1.**
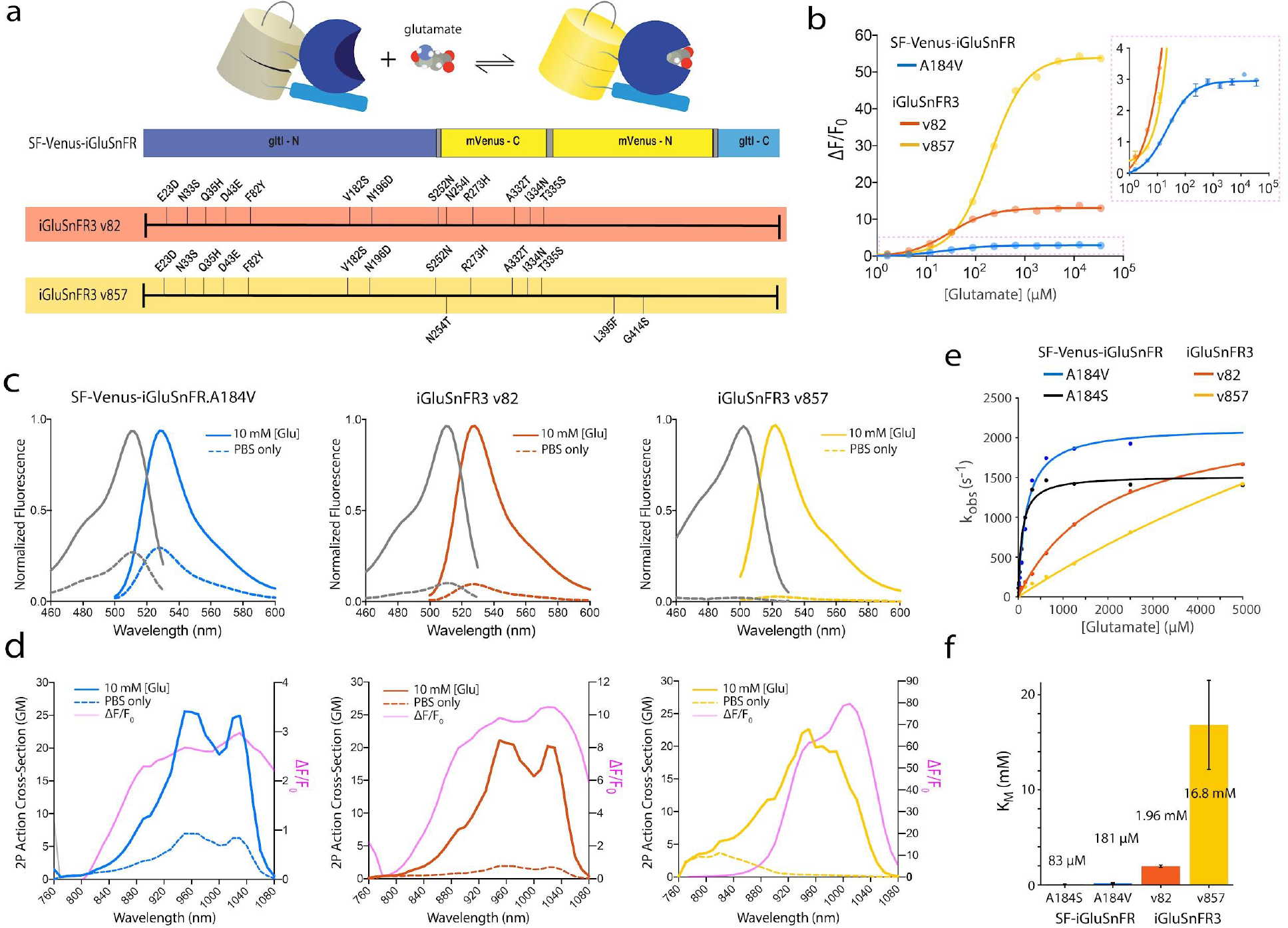
Photophysical properties of iGluSnFR3 variants. a) Top, iGluSnFR consists of a circularly-permuted fluorescent protein (yellow) inserted between interacting N- and C- terminal domains of the bacterial periplasmic glutamate/aspartate binding protein gltI (blue). Bottom, locations of amino acid substitutions of iGluSnFR3 variants v82 (red) and v857 (yellow) relative to the parent variant SF-Venus.iGluSnFR. b) Glutamate titrations of purified iGluSnFR protein. Right panel is a zoom-in view. C) 1-photon excitation and emission spectra of soluble SF-Venus-iGluSnFR.A184V, v82 and v857 in the presence (10 mM) and absence of glutamate. d) 2-photon excitation spectra and ΔF/F_0_ of the iGluSnFR3 variants in the presence (10 mM) and absence of glutamate. Pink lines are computed ΔF/F_0_. e) Stopped-flow measurements of initial ON rate (k_obs_) showing different degrees of kinetic saturation for SF-iGluSnFR and iGluSnFR3 variants. f) K_M_, the glutamate concentration that half-saturates the indicator’s initial ON rate. Responses to rapid changes in concentration above K_M_ are suppressed due to ON-rate saturation (Supplemental Note 1).

**Figure 2.**
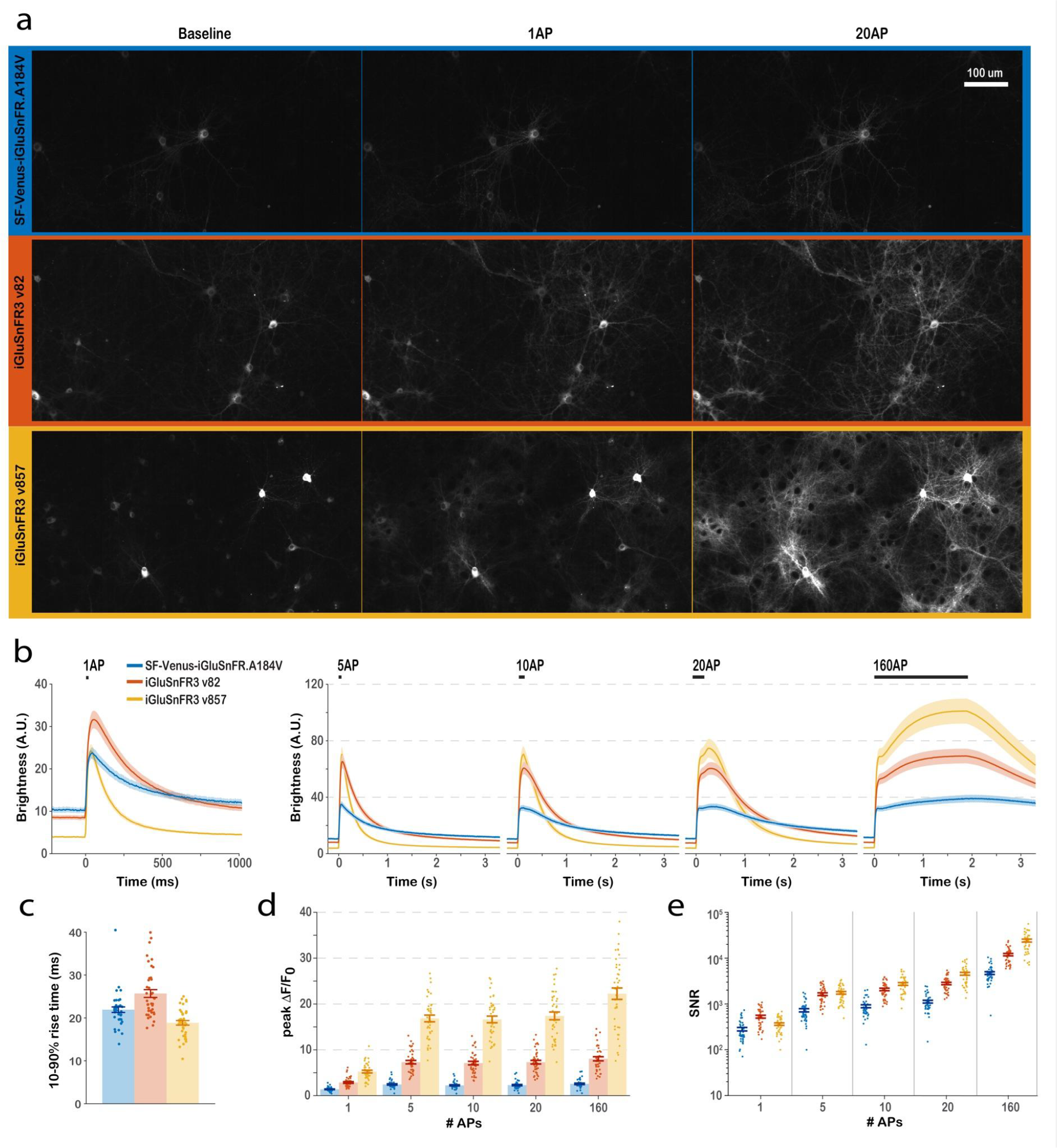
Characterization of iGluSnFR3 in cultured neurons. a) Images of primary cultures expressing surface-displayed SF-Venus-iGluSnFR-A184V (blue), v82 (red) and v857 (yellow) under hSyn promoter before stimulation and at peak brightness after field stimulations of 1 and 20 APs. b) Mean pixel brightness traces for stimulations of 1, 5, 10, 20, and 160 APs. c) 10-90% rise time of the three variants for the 1 AP condition. d) Peak ΔF/F_0_ of the three variants across conditions. e) Time-integrated signal-to-noise ratio for the three variants across conditions. Traces are mean ± SEM., N=40 culture wells for each variant.

As soluble proteins, v82 and v857 have less saturating activation kinetics than WT, with K_M_ values 1 and 2 orders of magnitude larger, respectively (WT: K_M_ = 180 ± 69 µM, v82: K_M_ = 1960 ± 210 µM, v857: K_M_ = 16800 ± 1550 µM; 95% c.i.; Supplemental Note 1; Figure 1d-e). Both show greatly increased dynamic range to saturating glutamate (1P Excitation ΔF/F_0_: {WT: 2.9 士 0.1; v82: 13.1 士 0.9; v857: 54.0 士 2.6} 2P Excitation ΔF/F_0_: {WT: 3.2 ± 0.5; v82: 10.5 ± 0.3; v857: 76.3 ± 13.5}; Figure 1b-c), due largely to dimmer glutamate-free states (Supplemental Table 1), and have high specificity to glutamate over other neurotransmitters, amino acids, and drugs (Supplemental Figure 1). v857 has an SYG rather than GYG chromophore sequence (Supplemental Note 3) and has a shifted fluorescence spectrum, larger glutamate-bound quantum yield (WT: Φ_SAT_ = 0.54 士 0.03, v82: Φ_SAT_ = 0.60 士 0.01, v857: Φ_SAT_ = 0.93 士 0.03), lower extinction coefficient (WT: ε_SAT_ = 116,000 士 3300 M^-1^cm^-1^, v82: ε_SAT_ = 105,000 士 1600 M^-1^cm^-1^, v857: ε_SAT_ = 40,000 士 2900 M^-1^cm^-1^), reduced pH sensitivity of the APO state (Supplemental Figure 2, Supplemental Note 2, Supplemental Table 1), and reduced soluble protein affinity (WT: K_d_ = 24 ± 2 µM, v82: K_d_ = 32 ± 4 µM, v857: K_d_ = 196 ± 15 µM) (Supplemental Figure 3, Supplemental Table 1). Despite a lower extinction coefficient, v857 has a large 2P action cross-section and exceptional 2P-excited molecular brightness measured by fluorescence correlation spectroscopy (Supplemental Table 1, Supplemental Figure 4). Unlike the other iGluSnFR variants, v857 exhibits strong quantum yield increase upon glutamate binding, which may be useful for generating probes with fluorescence lifetime contrast (25).

In cultured neurons, surface-displayed v82 and v857 express and traffic well (Figure 2a), have lower glutamate affinity than WT (WT: K_d_ = 2.0 ± 0.4 µM, v82: K_d_ = 4.5 ± 0.2 µM, v857: K_d_ = 8.2 ± 0.1 µM) and have larger responses to bath-applied glutamate (Supplemental Figure 5) and field-stimulated APs (Figure 2b,d). v82 has slower, and v857 has faster, 1 AP rise time than WT (Field-stimulated cultures at 21℃; WT: 21.9 士 0.6; v82: 25.7 士 0.9 ms; v857: 18.9 士 0.5 ms; Figure 2c). Although faster kinetics often result in lower time-integrated photon counts, v857 nevertheless produces higher time-integrated signal-to-noise ratio (SNR) than WT in all conditions tested, as does the slower v82 (Figure 2e).

Together, these results demonstrate that v857 is a sensitive and selective new variant of iGluSnFR with rapid on-neuron kinetics and a high K_M_, desirable for improving sensitivity to synaptic versus extrasynaptic signals. v82 is a sensitive and selective variant with higher affinity, slower on-neuron kinetics, and lower K_M_ than v857, properties that should make it comparatively more responsive to extrasynaptic glutamate.

### v857 shows increased sensitivity, linearity, and spatial specificity in culture

We tested the sensitivity, spatial specificity, and linearity of the iGluSnFR variants by monitoring signals caused by spontaneous and evoked glutamate release at synapses between cultured neurons. APs evoke synchronous release of varying numbers of vesicles per synapse. Additionally, ongoing spontaneous vesicle release produces asynchronous post-synaptic potentials (miniature release events) with amplitudes typically corresponding to single vesicles (26, 27). With APs silenced (1 μM TTX), synapses expressing v857 show large-amplitude asynchronous flashes of fluorescence (optical miniature release events, ‘optical minis’, (14, 15)). As with electrophysiologically-observed miniature release events (27), the rate of optical minis increases in hyperosmotic sucrose buffer (Supplemental Video 1, Figure 3a-c, Supplemental Figure 6). The large signals and high photostability of v857 allowed continuous recordings of optical minis for 15 minutes without reduction in SNR (Supplemental Figure 7). Despite exhibiting higher SNR for 1 AP field stimuli, v82 detected fewer optical minis than v857. WT detected still fewer minis than either iGluSnFR3 variant (Figure 3d). These differences demonstrate that large field stimulus responses do not necessarily predict detection of synaptic signals by iGluSnFR variants, and suggest that kinetic properties such as K_M_ and rise time may better correlate with synaptic signals.

**Figure 3.**
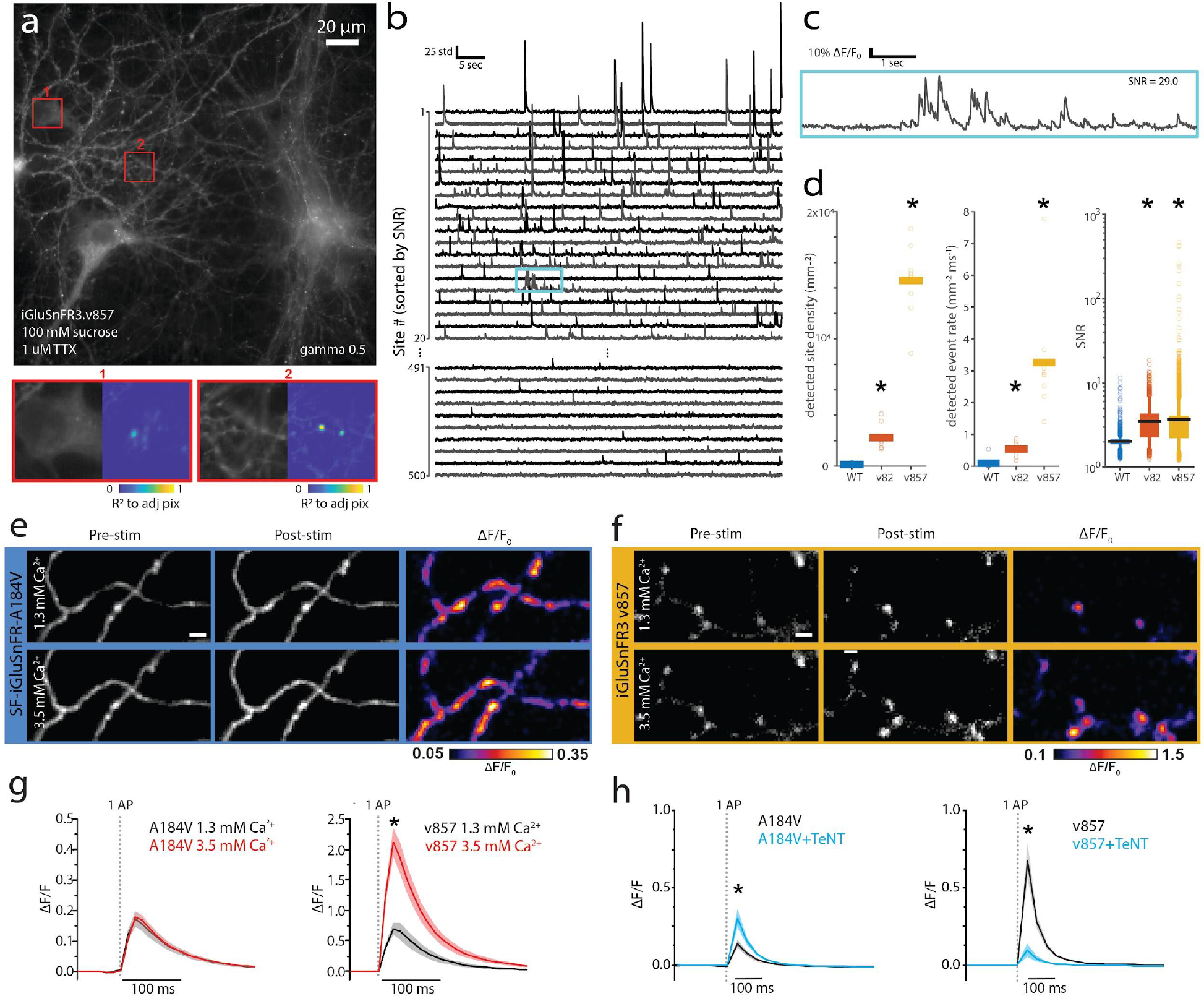
Recording and manipulation of synaptic release in primary neuron culture. a) (top) Image of representative example DIV 14 rat primary culture expressing v857. (bottom) Zoom in of regions highlighted in red squares above, accompanied by correlation images showing example sites of activity on a cell soma (1) and neurites (2). b) Activity traces for individual release sites in the same recording. Each trace is normalized to the standard deviation of noise at that site. Sites are sorted by SNR, with sites ranked 1-20 and 491-500 shown. c) Zoom-in of the trace highlighted in cyan in (b). d) Detected site density, event rate, and SNR across fields of view for WT and the two iGluSnFR3 variants expressed as hSyn.minDisplay constructs. N = 9 FOVs (v82 and v857); 8 FOVs (WT), from 3 culture wells per variant. (e-f) Representative images of SF-iGluSnFR.A184V (e) and v857 (f) in axons of sparsely-labeled cultured hippocampal neurons before and after 1 AP field stimulation, and corresponding ΔF/F0 images. Scale bar = 2 micrometers. g) Mean single-AP evoked fluorescence traces at boutons of cells expressing SF-iGluSnFR.A184V (left) and v857 (right) with bath calcium concentrations of 1.3 mM (black) and 3.5 mM (red). Dotted line denotes time of stimulus, shading denotes SEM. N=7 cells for each variant. *: p<0.01, paired Wilcoxon test. h) Mean single-AP evoked fluorescence traces at boutons of sparsely-labeled cells expressing SF-iGluSnFR.A184V (left) and v857 (right) co-transfected with tetanus toxin light chain (TeNT; cyan) or control (black). Dotted line denotes time of stimulus, shading denotes SEM. A184V control: N=6 cells, A184V+TeNT-LC: N=6 cells; v857: N=8 cells; v857+TeNT: N=7 cells. *: p<0.05, two-sample t-test.

To assess linearity of iGluSnFR variants, we imaged glutamate release evoked by single APs at individual axonal boutons. The probability of release, and thus the average number of vesicles released, depends nonlinearly on extracellular [Ca2+] (28, 29). Using v857, increasing extracellular [Ca^2+^] from 1.3 mM to 3.5 mM increased single-bouton 1AP ΔF/F responses from 70±9% to 213±24%, respectively (Figure 3e-g), consistent with previous measurements (28, 29). Under the same conditions, SF-iGluSnFR.A184V exhibited no significant increase in single-bouton response with increased extracellular calcium, consistent with kinetic saturation of that indicator.

To assess synaptic specificity of iGluSnFR variants, we compared field-stimulated 1 AP responses on axons in cultures in which a small subset of neurons were transfected with either iGluSnFR alone or with both iGluSnFR and the tetanus toxin light chain peptide (TeNT). TeNT cleaves synaptobrevin and blocks vesicle fusion (30) (Supplemental Figure 8). Axons of neurons coexpressing TeNT still showed responses, likely corresponding to extrasynaptic signals originating from nearby untransfected axons. The amplitude of these signals was greatly reduced in v857+TeNT neurons (Figure 3h) compared to responses of cells expressing v857 alone. In SF-iGluSnFR.A184V+TeNT co-expressing cells, these signals were no smaller than responses of cells expressing SF-iGluSnFR.A184V alone.

Together the above results indicate that v857 more linearly reports glutamate release at individual synapses and better distinguishes synaptic from extrasynaptic glutamate in neuronal culture, whereas SF-iGluSnFR.A184V undergoes saturation that limits its performance in these respects.

### Display constructs that improve optical mini signals and nanoscopic localization to post-synapses

The spatial specificity of glutamate transmission relies on nanoscopic localization of receptors to post-synaptic densities apposed to presynaptic active zones. However, electrophysiological and imaging studies have suggested that iGluSnFR may be excluded from the post-synaptic density, impacting the indicator’s ability to report synaptic transmission when expressed post-synaptically (13). We therefore compared a variety of surface display sequences for their performance at reporting synaptic signals, using the amplitude, SNR, and detection rate of optical minis as readouts.

The SnFR family of indicators are targeted to membranes via insertion between an N-terminal IgΚ secretion leader sequence and C-terminal PDGFR transmembrane domain (19). We compared v857 in this PDGFR construct to 15 C-terminal variants including GPI anchors, transmembrane domains, cytosolic motifs, and ER and Golgi export signals (Supplemental Table 2). Of these, two GPI anchors (GPI_COBL9_, GPI_NGR_) and a fusion to a modified cytosolic fragment of stargazin (SGZ) significantly increased the median SNR of optical minis (Figure 4a-b). These variants also exhibited significantly improved surface trafficking, as measured by ΔF/F upon bath application of glutamate (Supplemental Figure 9). A followup screen combining GPI_COBL9_ (henceforth, GPI) with 14 N-terminal leader sequences that we have used to optimize other cell-surface indicators (31) resulted in no further improvement compared to the existing IgK leader (Supplemental Figure 9).

**Figure 4.**
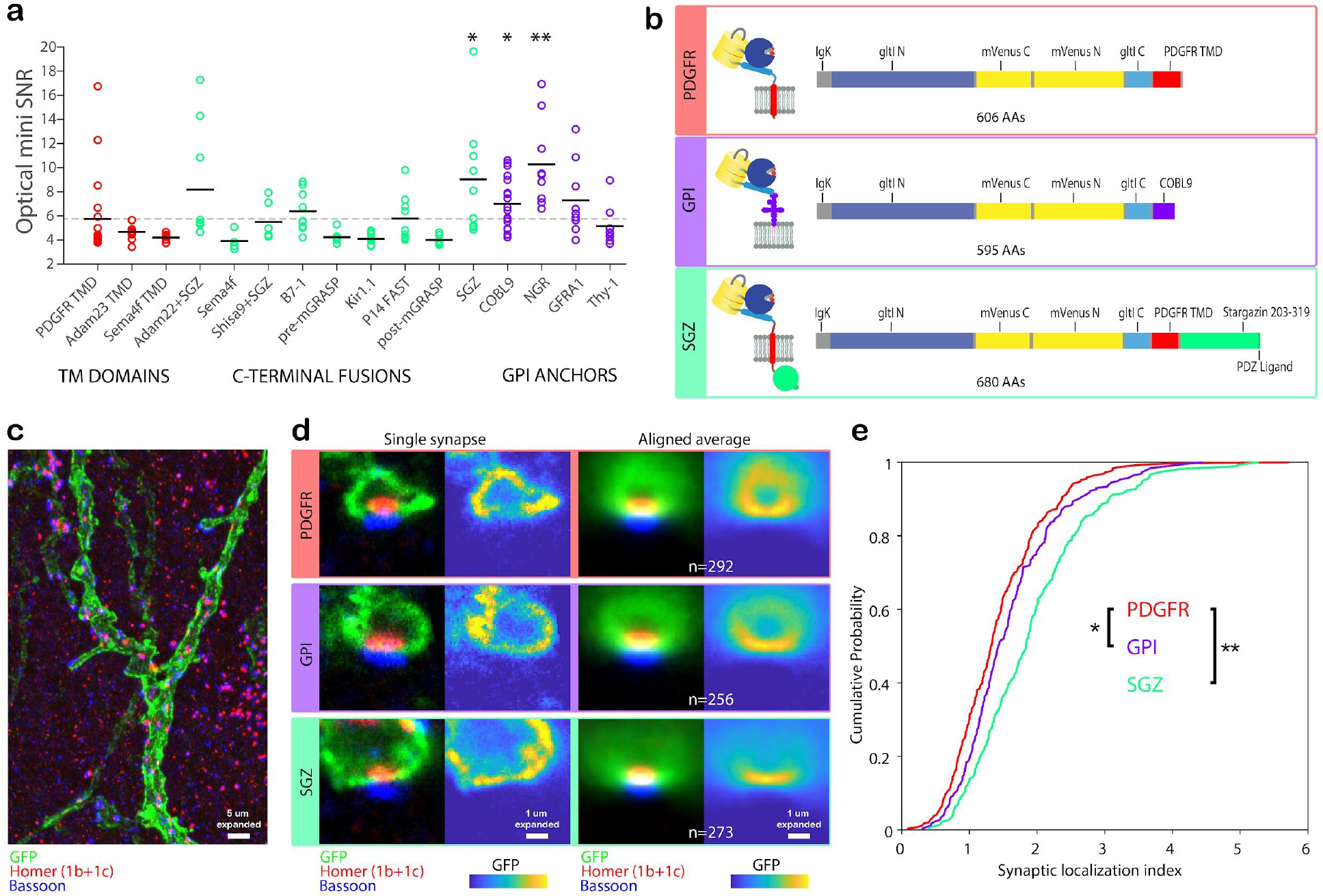
C-Terminal membrane anchoring sequences improve functional response and nanoscopic localization. a) SNR of optical minis in culture for v857, expressed using PDGFR transmembrane domain (pMinDisplay) and 15 other display constructs. For each construct, black line denotes the sample mean. N= 9 FOVs over 3 wells per construct. b) Sequence schematics for PDGFR, GPI, and SGZ display constructs. PDGFR is a C-terminal fusion to the PDGFR transmembrane domain in the mammalian expression pMinDisplay vector. GPI contains a C-terminal glycosylphosphatidylinositol anchor. SGZ contains the PDGFR transmembrane domain, followed by a modified form of the cytosolic C-terminal domain of Stargazin including a terminal PDZ ligand. c) Representative maximum intensity projection image of a portion of an expanded gel of mouse cortical tissue expressing v857.SGZ, immunostained for GFP (green), Homer1 (red), and Bassoon (blue). d) Images of single representative synapses and aligned averages for the three iGluSnFR variants. For each image, immunostains for GFP (green), Homer1 (red), and Bassoon (blue) are shown at left, and the GFP channel is shown at right. e) Cumulative probability plot of the ratio of GFP signal at the synapse center to that within a 3x6 µm (expanded) post-synaptic region. N = 292 (PDGFR), 256 (GPI), 273 (SGZ) synapses. *:p<0.05, **p<0.01 Kolmogorov-Smirnov test.

We next evaluated the *in vivo* nanoscopic localization of these functionally-improved constructs. We generated adeno-associated virus (AAV) encoding Cre-dependent SGZ, GPI and PDGFR variants of v857 and expressed them in the cortex of *Emx1*-Cre mice. We used expansion microscopy (32) to resolve post-synaptic densities with immunolabeling for GFP (iGluSnFR), a presynaptic (Bassoon) and a post-synaptic (Homer1) marker (Figure 4c). For all variants, anti-GFP labeling was heterogeneously distributed within the membrane, and gaps in labeling appeared throughout dendrites. However, reduced labeling of post-synapses specifically in PDGFR-labeled neurons was clearly observed by computing aligned average images over all identified synapses in expanded volumes (Figure 4d). Post-synaptic localization, *i.e.*, the ratio of mean anti-GFP signal at the synapse center to that within a 3x6 µm (expanded) post-synaptic region, was significantly increased for both GPI and SGZ constructs (PDGFR: 1.43±0.05; GPI: 1.57±0.05, p=0.037; SGZ: 1.78±0.04, p<1e-10; two-sample Kolmogorov-Smirnov tests against PDGFR; Figure 4e).

Together these results identified motifs that improve expression and trafficking of iGluSnFR in cultured neurons, increase the SNR of synaptic glutamate signals, and improve the indicator’s post-synaptic localization in vivo.

### In vivo iGluSnFR3 imaging and electrophysiology in visual cortex

We next characterized iGluSnFR3 in vivo in the visual cortex of mice. We observed strong signals in mice injected with AAVs encoding v857.GPI or v857.PDGFR variants, including large-amplitude responses to visual stimuli in 1-photon widefield recordings (Supplemental Figure 10), large-amplitude transients localized to spines and boutons in 2-photon recordings (Supplemental Video 2), and greatly increased photostability relative to WT (Supplemental Figure 11). Although immunohistological labeling of virally-expressed SGZ indicated strong expression and good membrane localization in cortex in vivo (Figure 4c-d), and we observed bright viral expression and functional signals in rat cortical culture (not shown), SGZ exhibited dimmer native fluorescence and weaker functional responses than PDGFR and GPI variants in mouse cortex (Supplemental Figure 10). We therefore performed all remaining in vivo experiments with the PDGFR and GPI variants.

Glutamate imaging is a promising approach for studying spatiotemporal patterns of synaptic activity in the intact brain (10,18,20,33) but the relationship between in vivo iGluSnFR signals and synaptic activity, such as presynaptic APs, has not been characterized. To characterize this relationship, we first imaged individual boutons on axons of individual sparsely-labeled glutamatergic neurons during simultaneous cell-attached electrophysiological recordings from the parent cell bodies. Layer 2/3 pyramidal neurons in the mouse primary visual cortex were electroporated with DNA plasmids encoding either v857.GPI or SF-iGluSnFR.A184S. Cell-attached recordings and concurrent 2P imaging at 500 Hz frame rate were performed in lightly anesthetized, head-fixed mice (Figure 5a,b).

**Figure 5.**
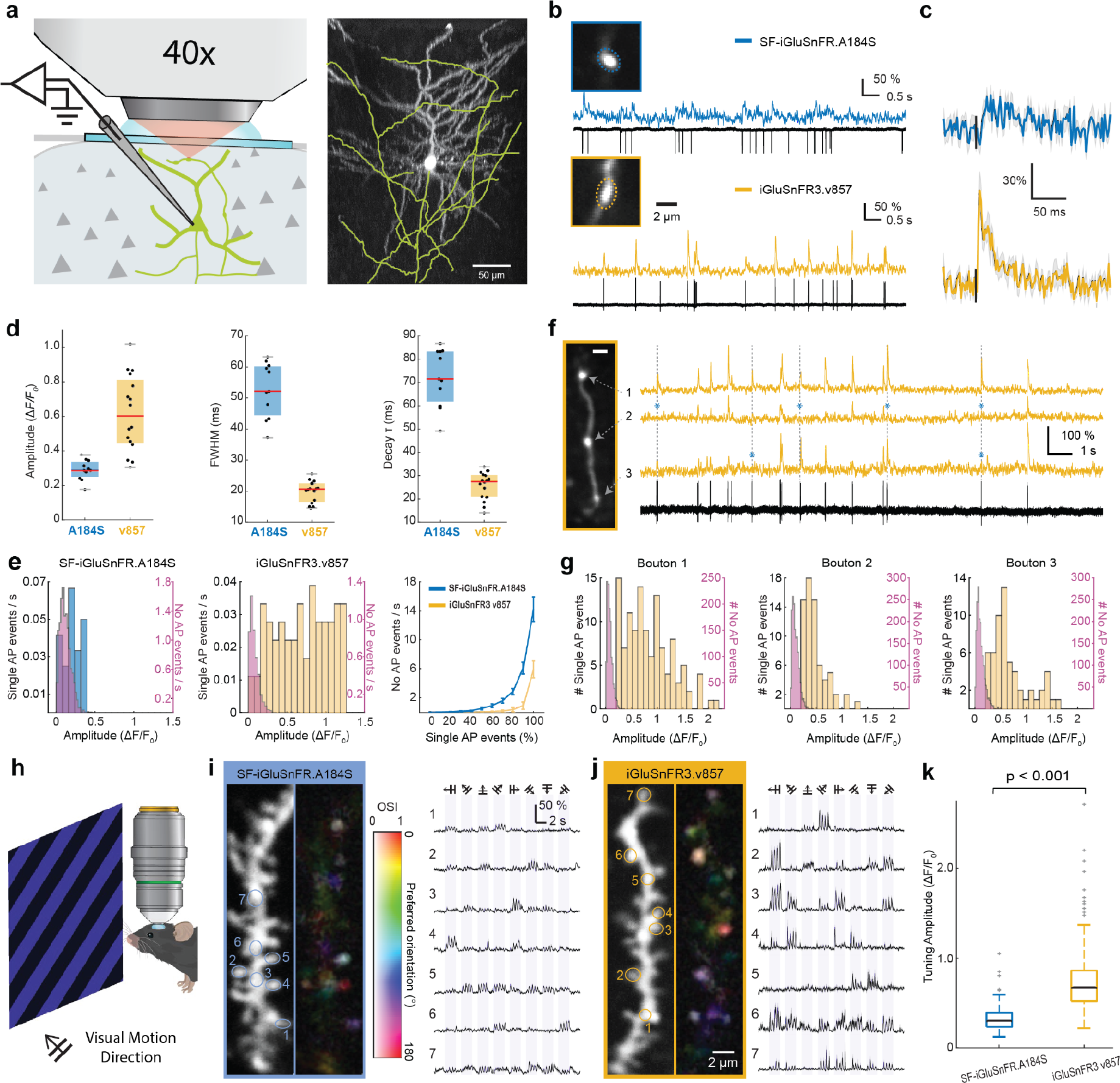
In vivo two-photon glutamate imaging and electrophysiology in visual cortex. a) (left) Experimental scheme illustrating simultaneous two-photon axonal imaging and cell-attached recording in vivo. (right) Z-projected image of a iGluSnFR3.v857 labeled neuron in mouse V1. The axon is traced and shown in yellow. b) Two-photon imaging of individual boutons and simultaneous cell-attached recordings from representative example neurons labeled with SF-iGluSnFR.A184S (top, “A184S”) and v857 (bottom). c) Mean fluorescence signal for isolated action potentials measured with v857 (bottom) and A184S (top). Each trace is an average of n=10 consecutive events. d) Amplitude and kinetics of fluorescence signals for isolated action potentials measured with v857 (n=16 boutons from 5 neurons) and A184S (n=11 boutons from 5 neurons). e) Left and middle panels: histograms of amplitudes for single-AP events and in absence of APs for example boutons shown in (b). Right panel: comparison of the non-AP event rate for v857 (yellow) and A184S (blue) as a function of detection threshold (mean ± SEM, n=6 boutons for v857 and n=5 boutons for A184S). f) Simultaneous v857 imaging (yellow) and cell-attached electrophysiological recording for three neighboring boutons recorded at 500 Hz each. Asterisks indicate synaptic release failures. Scale bar 2 um. g) Distribution of deconvolved AP-associated event amplitudes for the three boutons shown in (f). h) Experimental schematic for in vivo imaging of responses to visual motion stimuli in dendrites in primary visual cortex (V1). i) Left, two-photon image of a dendritic segment of a cortical L2/3 neuron expressing A184S, and the corresponding pixel-wise orientation tuning map (hue: preferred direction, saturation: OSI, lightness: response magnitude). Right, single-spine glutamate transients evoked by drifting grating visual stimulations for the directions indicated on top. Numbers correspond to spines indicated in image at left. j) Same as (i) but from a L2/3 neuron expressing v857 in a different mouse. Color scales for the tuning map are identical to those in (i). k) Comparison of glutamate transient amplitudes to visual stimulations for v857 and A184S (0.6140 [0.4620-0.8162] vs 0.2944 [0.2164-0.3925], p<0.001, n=261 ROIs from 5 neurons for v857 and n=375 ROIs from 5 neurons for A184S). Data are expressed as median [Interquartile range].

Boutons of v857-expressing neurons exhibited larger and faster 1AP-associated transients than those expressing SF-iGluSnFR.A184S (Figure 5c,d). To quantify the specificity with which fluorescence transients report synaptic release, we compared the distribution of transient amplitudes associated with single electrophysiologically-observed APs to those not associated with APs. v857 showed a greater difference between AP-associated transients and non-AP-associated events than A184S (Figure 5e). Using an amplitude threshold on denoised traces, 70% of presynaptic APs were detected with v857 at a false positive rate of 0.2 Hz, while 70% of presynaptic APs were detected with SF-iGluSnFR.A184S at a false positive rate of 3.7 Hz (Figure 5f). At this threshold, we attribute most of the APs undetected by v857 to failures of synaptic release, since imaging measurement noise was far lower than the variability in single-AP event amplitudes. As a demonstration, we imaged three neighboring boutons on a single axon during cell-attached recording of APs from the parent cell body, and summarized the amplitude distribution of AP-associated fluorescence transients collected from each bouton during 1.5 min of continuous recording (Figure 5f,g). We next recorded from labeled dendrites while presenting visual motion stimuli to lightly-anesthetized mice (Figure 5h). v857 reported direction-tuned responses that were more localized to dendritic spines and had significantly larger amplitudes than those reported by SF-iGluSnFR.A184S (Figure 5i,j,k).

### In vivo imaging of ascending and recurrent inputs to barrel cortex L4

We next used iGluSnFR3 to study feedforward and recurrent inputs to vS1, the primary vibrissa (whisker) sensory cortex. Rodents rhythmically sweep their vibrissae back and forth to sense objects through touch (34). Touch and self-motion signals ascend from the vibrissal follicles through ventral posterior (VPM) thalamus, then to L4 of vS1. Each vibrissa is represented by a column of neurons in vS1 forming a characteristic “barrel” structure. L4 excitatory cells within a barrel receive about 10% of their excitatory input from thalamocortical (TC) projections (35), with most remaining (recurrent) connections originating from other L4 excitatory cells in the barrel (36, 37). The depth of L4 below the brain surface (∼350-450 µm) and the small volume of individual synapses (<1 femtoliter) are challenges for measuring precisely-timed synaptic activity. To overcome these challenges, we took advantage of iGluSnFR3’s high SNR and used direct wavefront sensing adaptive optics two-photon (AO2P) microscopy to achieve diffraction-limited spatial resolution (33).

We used two different strategies to label TC versus recurrent synaptic transmission (Figure 6a). To express iGluSnFR variants in thalamocortical axons, AAV was injected into VPM. Anatomically-defined thalamic boutons in L4 of awake, head-fixed mice showed large-amplitude fluorescence transients induced by vibrissal touch during rhythmic active whisking (Supplemental Figure 12). To quantify TC responses across frequencies, we used rhythmic passive air-puff stimulation (Figure 6b) in awake mice, and characterized all detectable responding sites (Figure 6c). Rhythmic stimulation from 2 to 30 Hz produced time-locked rapid-onset responses (Figure 6d, Supplemental Video 3) that were characterized for five iGluSnFR variants as a function of stimulation frequency (Figure 6e; 1609 individual boutons across 13 mice; Supplemental Table 3 lists variants used). v857 showed greater modulation depth than the SF-iGluSnFR variants at all stimulation frequencies, over 2-fold greater than the green variant SF.iGluSnFR.A184V and 5-fold greater than SF-Venus.iGluSnFR.A184V for 2 to 20 Hz stimulation rates. Using v857, our AO2P microscope sampling at 125 Hz was able to resolve single-trial responses at rates of 15 Hz or slower with a signal-to-noise ratio above 1.0.

**Figure 6.**
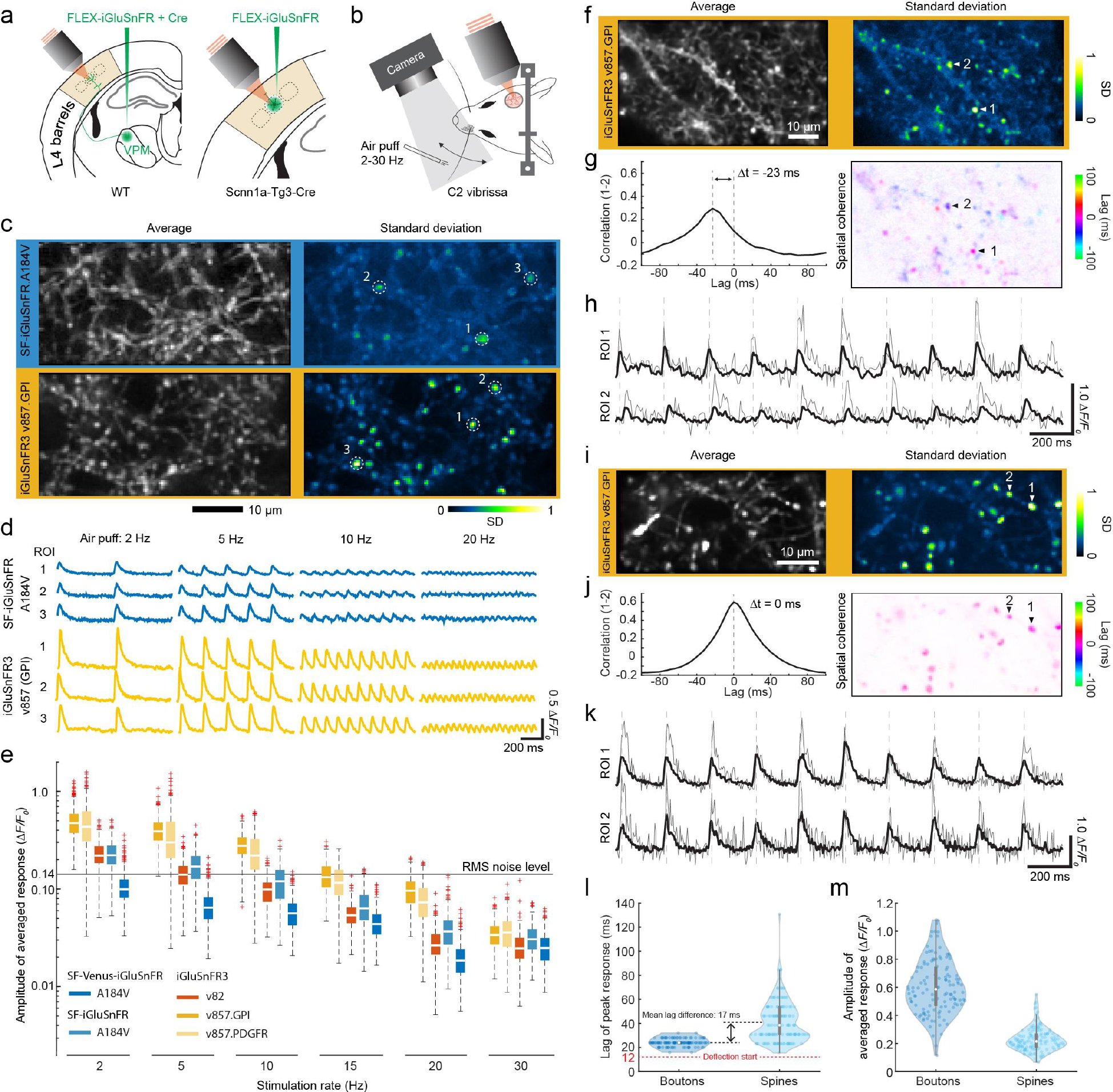
Adaptive optics glutamate imaging of thalamocortical boutons and dendritic spines in Layer 4 of vibrissa sensory cortex. a) Schematic of AAV injection in VPM (left) in wild-type mice to express iGluSnFR variants on thalamocortical boutons and, separately, injection into vibrissa cortex (right) in *Scnn1a*-Tg3-Cre mice to express iGluSnFR on dendritic spines of L4 cortical neurons. b) Schematic of air-puff vibrissal stimulation in awake mice during high-speed adaptive optics 2P imaging. Frame rates were ∼250 Hz for boutons (c-e, i-k) and ∼130 Hz for spines (f-h). c-e) Comparison of iGluSnFR3 and SF-iGluSnFR variants in TC boutons for different stimulation frequencies. c) Left: average image of TC axons and boutons in L4 vibrissal cortex labeled by SF-iGluSnFR.A184V (top) and v857.GPI (bottom). Right: normalized pixel-wise standard deviation (SD) across 1-s averaged epochs from 60 trials. C) Mean responses across 60 trials for representative boutons shown in panel c labeled by v857.GPI (top) and A184V (bottom). e) Comparison of averaged response amplitudes of TC boutons labeled with iGluSnFR3 and SF-iGluSnFR variants: v857.GPI (3 mice, 389 boutons), v857.PDGFR (3 mice, 314 boutons), v82.GPI (2 mice, 312 boutons), A184V (3 mice, 239 boutons) and Venus-A184V (2 mice, 355 boutons). Line indicates the root-mean-square (RMS) noise level calculated from trials for v857.GPI. f-h) Imaging of dendritic spines labeled with v857.GPI to resolve response lag of different L4 spines to vibrissal stimulation at 5 Hz. f) Mean (left) and normalized SD (right) images of dendritic spines in L4 vibrissal cortex labeled by v857.GPI. g) Left: cross-correlation of fluorescence signals from two selected active spines highlighted at right. Right: pixelwise lag of peak response relative to the dominant mode in the recording. h) Time series from the same two dendritic spines in panel g with a representative trial in gray and the mean over 10 trials in black. i-k) Imaging of TC boutons labeled by v857.GPI analyzed and displayed as in (f-h). I) Comparison of lags of the peak response for TC boutons (123 ROIs) and dendritic spines (145 ROIs) in L4 vibrissal cortex. Time 0 denotes onset of electronic trigger to the air-puff valve, while the red dashed line indicates measured onset of vibrissal deflection. Violin plots. m) Mean response amplitudes of TC boutons (123 ROIs) and dendritic spines (145 ROIs), violin plots.

The high signal-to-noise ratio and synaptic specificity of v857 permitted us to map the temporal lags of TC and recurrent inputs. To record from L4 dendrites we injected AAV encoding Cre-dependent v857.GPI into vS1 of *Scnn1a*-Cre mice, which express Cre recombinase in L4 excitatory neurons (38). v857.GPI showed equally strong labeling of cortical spines as with TC boutons (Figures 6c,f). We recorded spine responses, which should predominantly reflect recurrent input, while stimulating the vibrissae of awake mice at 5 Hz (Supplemental Video 4). Spine responses each showed a characteristic lag that differed across pairs of synapses by up 70 ms as revealed by space-frequency analysis (39) (Figure 6g) and by directly comparing time-averaged responses (Figure 6h). Similar comparisons across 123 TC boutons from 2 mice showed no differences in time lags (Figure 6i-k). As a population (145 dendritic spines from 2 mice) the lags of recurrent connections are broadly distributed with an average delay within layer 4 alone of 17 ms (Figure 6l). The absence of a heterogeneous set of time-lags for the dense network of axons imaged in VPM-labeled mice further demonstrates the high specificity of v857 for synaptic versus extrasynaptic signals in vivo.

## Discussion

Synaptic transmission achieves high spatial specificity in large part from the kinetics and localization of endogenous glutamate receptors. Motivated by these biophysical principles, we engineered a glutamate indicator with high spatial specificity at synapses and greatly improved in vitro and in vivo performance. iGluSnFR3 achieves this performance through its rapid kinetics and high rate-limiting glutamate concentration (K_M_) that improve indicator linearity and spatial specificity, and shows improved nanoscopic localization to post-synapses via the GPI and SGZ C-terminal membrane-anchoring sequences.

Glutamate imaging has long promised to complement or replace dendritic Ca^2+^ imaging for recording spatially-resolved synaptic input patterns. The two modalities report different aspects of synaptic transmission and present distinct advantages and confounds. Dendritic Ca^2+^ imaging reports post-synaptic signals that are governed largely by (nonlinear) NMDA receptor activation in spines (40), but are less-studied at non-spiny synapses, and are spatially mixed by nonlocal processes such as backpropagating potentials and dendritic spikes (7). Glutamate imaging avoids these confounds, providing advantages for studying spatial arrangements of inputs and their correlations. However, functional imaging of membrane-bound indicators poses challenges compared to cytosolic ones (41), demanding indicators with exceptional sensitivity and photostability. Furthermore, accurate measurement of synaptic inputs requires linearity and spatial specificity. iGluSnFR3 was engineered to improve all of these properties to facilitate synaptic input measurements.

iGluSnFR3 has enabled studies that were previously infeasible. Firstly, we demonstrated the improved linearity of v857 in multiple contexts including purified protein (Figure 1e,f), field stimulation (Figure 2d), and individual synapses in culture (Figure 3g) and in vivo (Figure 6e). This enables more quantitative measurements of glutamate release over space and time that were previously confounded by saturation effects. Similar advantages were recently demonstrated for another novel variant in culture (13). Secondly, we demonstrated reliable detection of glutamate release associated with electrophysiologically-identified APs in vivo (Figure 5a-g). Such recordings provide ground-truth data for assessing the sensitivity of indicators and validating their synaptic specificity, and will provide quantitative constraints for the interpretation of signals in synaptic imaging studies. Lastly, we have now achieved sufficient SNR, kinetics, and spatial specificity to record from populations of synapses while resolving differences of a few milliseconds in the timing of individual inputs. These capabilities will be valuable for studying the synaptic dynamics underlying neuronal computation and learning (9,11,42).

## Methods

### Animal care and use statement

All experimental procedures involving animals were performed in accordance with protocols approved by the Institutional Animal Care and Use Committee at the respective institute ((HHMI Janelia Research Campus, Dartmouth College, University of California San Diego, and TUM). Procedures in the USA conform to the NIH Guide for the Care and Use of Laboratory Animals. Procedures at TUM were approved by the state government of Bavaria, Germany .

HHMI Janelia Research Campus: protocols 14-115, 16-225 Dartmouth College: protocol 00002115 University of California, San Diego: protocol S02174M Technical University of Munich: ROB-55.2-2532.Vet_02-17-181 This research has complied with all applicable ethical regulations.

### Molecular biology

Site-directed mutagenesis was performed using QuikChange (Agilent Technologies) or Q5-site directed mutagenesis kit (New England Biolabs) according to the manufacturer’s suggestions. Error-prone PCR was done using Mutazyme PCR from Genemorph II mutagenesis kit (Agilent Technologies) according to the manufacturer’s suggestions, generating mean mutation rates of 4.5 to 9 DNA substitutions/clone. Recombination libraries were made using Staggered Extension PCR (StEP). In each case, mutagenesis was performed on the coding sequence and reinserted into the same plasmid vector via Gibson assembly or digestion-ligation. Plasmid sequences were confirmed via Sanger sequencing.

### Construction of mammalian expression vectors (Janelia)

To express sensors on the cell surface, iGluSnFR coding sequences were PCR amplified from the bacterial expression (pRSET) constructs, followed by Gibson assembly into pAAV.hSyn (or Cre-dependent pAAV.hSyn.FLEX) vectors, into which the N- and C- terminal flanking sequences (IgK leader and surface display sequence) were first inserted. For the development of the display constructs library, the N-terminal leader or C-terminal surface display sequence was substituted by Gibson assembly. Recombinant AAV particles (rAAVs) were prepared by Viral Tools at Janelia. Briefly, 293T cells were prepared in 3 of 150 mm T/C dishes at 1E7 cells/dish 2 days before transfection. Cells were transiently transfected with a total of 84 mg of DNA at a ratio of pHelper plasmid: capsid plasmid: AAV construct = 3:2:5 using polyethylenimine. The transfected cells were replenished with fresh serum-free clear DMEM at 6∼8 hours post-transfection and incubated for 3 days in a CO2 incubator (37 °C and 5% CO2). rAAVs were collected from both cells and supernatant and purified by two rounds of continuous cesium chloride density gradient. The rAAV preps were dialyzed, concentrated to 100 uL, and sterilized by filtration. The final viral titers were measured by qPCR on the viruses’ ITR.

### Construction of display construct library

Synthetic DNA consisting of different N-terminal secretion peptides and C-terminal anchoring domains were codon-optimized for *Mus musculus* and ordered as gene blocks from Integrated DNA Technologies (IDT). The fragments were PCR amplified and inserted into a pAAV.hSyn iGluSnFR vector via Gibson assembly. For the insertion of N-terminal secretion peptides, the vector backbone was created by digesting the pAAV.hSyn.iGluSnFR gene with NheI and BamH1 to cut out IgK sequence and substituting the new fragments. For C-terminal anchors, the iGluSnFR gene was digested with PstI and HindIII, to cut out PDGFR and substituting the other anchors. The C-terminal libraries were cloned in combination with IgK as N-terminal secretion peptide, and N-terminal secretion peptide libraries were cloned with GPI_COBL9_ as the C-terminal anchoring domain.

### Bacterial colony and lysate measurements

iGluSnFR variants in soluble form in pRSET vector (Thermo Fisher Scientific) were transformed via electroporation into competent *E. coli* bacteria and plated. Fluorescence was excited with 488 nm light on an LED light table (for 1-photon screening) or 1030 nm using a custom laser scanning system (for 2-photon screening). The 2-photon screening system used a 56-Watt, 160 fs, 1MHz repetition rate fiber laser (Tangerine HP2, Amplitude Systemes), expanded into a 20 mm long line focus and scanned with a line scanner to illuminate a ∼2x4 cm field of view (Supplemental Note 2). In each round, 400 to 1000 highly fluorescent colonies were picked, and isolated into 96-Well Polypropylene DeepWell blocks (Thermo Fisher Scientific) containing 800 µL of Studier’s autoinduction medium. The DeepWell cultures were grown at 30 °C, 250 rpm for 30 hours before cells were harvested by centrifugation (8,000 g for 10 min), and rapidly lysed using liquid nitrogen and 40 °C water bath freeze-thaw cycles. 100 µL of resulting crude protein extract was transferred to a glass-bottomed plate, in which fluorescence spectra were collected before and after addition of 10 µL of glutamate buffer in PBS (137 mM NaCl, 2.7 mM KCl, 10 mM Na_2_HPO_4_, 1.8 mM KH_2_PO_4_; pH 7.3), using a plate reader (Tecan Infinity M1000). The final glutamate concentration used varied over rounds (200 µM - 2 mM) as the affinity of variants under test changed, but were consistent within any given round. In some rounds, multiple final glutamate concentrations were used for lysate measurements to estimate both affinity and dynamic range.

### Protein purification and *in vitro* characterization

pRSET bacterial expression vector (Thermo Fisher Scientific) encoding the soluble form of the iGluSnFR protein of interest was chemically transformed into *E. coli* strain T7 Express Competent *E. coli* (New England Biolabs). A 4 mL culture containing LB + 100 µg/mL ampicillin, inoculated with a single bacterial colony, was grown overnight (37 °C, 250 rpm) before being diluted into 500 mL Studier’s autoinduction medium^2^. The culture was grown at 30 °C, 250 rpm for 30 hours before cells were harvested by centrifugation (8,000 g for 10 min). The resulting lysate was purified using Ni-NTA chromatography using HisPur Ni-NTA beads (Thermo Fisher Scientific), and dialyzed using Slide-A-Lyzer dialysis cassettes (Thermo Fisher Scientific) with PBS for 24 hours. Glutamate-saturated and glutamate-free measurements were performed in 100 mM glutamate in PBS and PBS only, respectively. Glutamate titrations were performed using a series of serially diluted buffers. The fluorescence measurements were fit with a Hill equation: 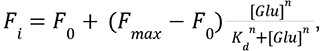, where F_0_ and F_max_ are fluorescence without glutamate and with saturated glutamate, respectively, and n is the Hill coefficient.

Observed apparent Hill coefficients were in all cases 1 ± 0.2. pH titrations were performed by making a series of 100 mM glutamate in PBS buffers and PBS only buffers with varying pH ranging from 5 to 9 (pH adjusted using concentrated HCl and NaOH). Fluorescence intensities as a function of pH were measured in both glutamate-saturated and glutamate-free states, and fitted with a sigmoidal binding function to determine the apparent p*K*a and apparent ΔF/F as a function of pH. The sigmoidal binding function used was fit using Graphpad Prism 8 software: 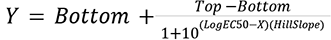, where Top and Bottom are limit brightnesses at high and low pH, respectively, HillSlope is the steepness of the curve, and EC50 is the apparent pKa. Apparent ΔF/F_0_ was computed using the raw fluorescence (Y) values of glutamate-bound and glutamate-free state with respect to pH’s (X) between 5 and 9: 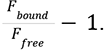. Specificity measurements were performed by making buffered solutions in PBS of 20 conical L-amino acids at 100 mM (pH 7.3), neurotransmitters at 100 mM (pH 7.3), and other drugs at varying concentrations: DNQX (10 µM), D-AP5 (5 mM), DL-TBOA (5 mM), NBQX (5 mM), NMDA (10 mM), CNQX (1 mM), kainate (5 mM).

For L-amino acids and neurotransmitters, single point excitation intensity (Ex: 510 nm, Em: 540 nm) was obtained in glutamate-free (APO) condition. ΔF/F was computed by using the following equation: (average fluorescence intensity with L-amino acid/neurotransmitter)/(average fluorescence intensity in apo)-1. For various drugs, single point excitation intensity (Ex: 510 nm, Em: 540 nm) was obtained in both APO and glutamate-bound (SAT) conditions. ΔF/F for each drug was independently computed in both APO and SAT conditions using the following equation: (average fluorescence intensity with drug)/(average fluorescence intensity in apo or sat)-1.

### One-photon photophysical measurements (Janelia)

1-photon fluorescence intensity spectra in presence and absence of glutamate were obtained using the Tecan Infinity M1000 plate reader. Excitation scans were performed from 460 nm to 530 nm (emission set at 550 nm, bandwidth 10 nm, step 2 nm), emission scans were performed from 500 nm to 600 nm (excitation set at 480 nm, bandwidth 10 nm, step 2 nm). Single point excitation intensities for glutamate titrations were obtained by exciting at 510 nm and collecting at 530 nm (bandwidth 10 nm). Extinction coefficient was determined using the alkali denaturation method, determining the denatured chromophore concentration using the extinction coefficient of denatured GFP as a reference (44,000 M^-1^ cm^-1^ at 447 nm). The extinction coefficient was computed by dividing the absorbance in non-denatured state by the calculated chromophore concentration. Quantum yield measurements were performed using a Quantaurus-QY spectrophotometer (Hamamatsu Photonics, Hamamatsu City, Japan), for purified proteins in PBS and 20 mM glutamate.

### Stopped-flow kinetics (Janelia)

Measurements of ON kinetics were made using an applied Photophysics SX20 Stopped-flow Reaction Analyzer using fluorescence detection, exciting at 505 nm and detecting with a 520 nm long pass filter, mixing equal volumes of dilute purified indicator protein and PBS+glutamate buffer. Measurements were performed at room temperature (22°C). Recorded intensity traces were fit in Matlab; code is available in the code and data package. At least 3 replicates per concentration were averaged, and a monoexponential saturation curve was fit to the average trace over the time window [t_25_ 4×t_90_], where t_25_ is the time at which intensity increased to 25% of maximum or 2 ms, whichever was sooner, and t_90_ is the time at which intensity increased to 90% of maximum. The rate-concentration relationship was then fit to the Michaelis-Menten model using Matlab’s *fit* function. Errors of fit were computed from the *fit* objects using the Matlab built-in *confint*.

### Two-photon photophysical measurements (Janelia)

2P measurements were performed on protein solutions (2-4 µM) in PBS with 0 mM (apo) and 100 mM (sat) added glutamate, using an inverted microscope (IX81, Olympus) equipped with a 60x, 1.2 NA water immersion objective (Olympus). Excitation was performed with an 80 Mhz Ti-Sapphire laser (Chameleon Ultra II, Coherent) to collect 2P excitation spectra from 710 to 1,080 nm. Fluorescence collected by the objective was passed through a shortpass filter (720SP, Semrock) and a band pass filter (550BP200, Semrock), and detected by a fiber-coupled Avalanche Photodiode (APD) (SPCM_AQRH-14, Perkin Elmer). The brightness spectra were normalized for 1 μM concentration and further used to obtain action cross-section spectra (AXS) with fluorescein as a reference. Fluorescence correlation spectroscopy (FCS) was used to obtain the 2P molecular brightness of the iGluSnFR3 soluble proteins at 950 nm and 1030 nm. The peak molecular brightness was defined by the rate of fluorescence obtained per total number of emitting molecules. 50-200 nM soluble iGluSnFR protein solutions were prepared in 100 mM glutamate PBS buffer and excited with 960 nm and 1030 nm wavelength at powers ranging from 2-30 mW for 200 seconds. The obtained fluorescence was collected by an APD and fed to an autocorrelator (Flex03LQ, Correlator.com). The obtained autocorrelation data were fit to a diffusion model to determine the number of molecules <N> present in the focal volume. The 2-photon molecular brightness (*ε*) at each laser power was calculated as the average rate of fluorescence *<F>* per emitting molecule *<N>*, defined as *ε* = *<F>/<N>* in kilocounts per second per molecule (kcpsm).

### Preparation of neuronal culture (Janelia)

Neonatal rat pups (P0) were euthanized; cortical and hippocampal tissue was dissected, and dissociated in papain enzyme (Worthington Biochemicals, ∼35 U/cortical and hippocampal pair) in neural dissection solution (10 mM HEPES pH 7.4 in Hanks’ Balance Salt Solution) for 30 minutes at 37 °C. After 30 minutes, enzyme solution was aspirated out and tissue pieces were subjected to trituration in 10% fetal bovine serum containing MEM media. Following trituration, cell suspension was filtered through a 40 μm strainer, and resulting single cell suspension was centrifuged. Cell pellet was resuspended in plating media (28mM glucose, 2.4mM NaHCO3, 100ug/ml transferrin, 25ug/ml insulin, 2mM L-glutamine, 10% fetal bovine serum in MEM) and cell counts were taken. Electroporation was conducted using the Lonza/Amaxa 4D nucleofector according to the manufacturer’s instructions, using 500 ng of plasmid and 5x10^5^ viable cells per transfection. Transfected cells were then seeded into 3 replicate wells in poly-D-lysine-coated glass-bottom 24-well plates or 35 mm dishes, and cultured at 37 °C with 5 % CO_2_. Cultures were fed twice a week by replacing 50% of the medium with fresh NbActiv.

### Glutamate titrations in neuronal culture (Janelia)

Experiments were performed with rat hippocampal and/or cortical primary cultures from neonatal (P0) pups. 14 days post transfection, culture medium was exchanged with 1 mL imaging buffer solution (145 mM NaCl, 2.5 mM KCl, 10 mM glucose, 10 mM HEPES, 2 mM CaCl_2_, 1 mM MgCl_2_, pH 7.3). Glutamate-free apo widefield images were taken at the center of each well using a Nikon Eclipse microscope (20X 0.4 NA, 505 nm excitation, 525/50 emission). Glutamate-saturated widefield images were taken at the same field of view (FOV) after the addition of 100 µL of 100 mM extracellular glutamate in imaging buffer (100 mM glutamate, 145 mM NaCl, 2.5 mM KCl, 10 mM glucose, 10 mM HEPES, 2 mM CaCl_2_, 1 mM MgCl_2_, pH 7.3). On-membrane glutamate titrations were performed by exchanging the culture medium with 1 mL of imaging buffer solution containing varying levels of glutamate concentration. A scalar constant background was first subtracted from all images. We quantified full-field ΔF/F for each FOV as: (mean brightness of SAT image)/(mean brightness of APO image)-1.

### Imaging optical minis (Janelia)

iGluSnFR3 variants were transfected in mixed hippocampal and neocortical primary cultures from neonatal (P0) rat pups. 14 days post transfection, culture medium was exchanged with 1 mL hyperosmotic buffer containing sucrose (100 mM sucrose, 145 mM NaCl, 2.5 mM KCl, 10 mM glucose, 10 mM HEPES, 2 mM CaCl_2_, 1 mM MgCl_2_, pH 7.3). 2 µM of tetrodotoxin citrate (TTX; diluted in deionized water) was added to each culture medium to silence the firing of action potentials. Videos were recorded for 60 seconds at 100 frames/s across 3 fields of view per well using a Nikon Eclipse microscope (60X 0.4 NA, 505 nm excitation center wavelength, 525/50 nm emission band). Additional experiments (Supplemental Figures 4,5) were performed with a spinning disk confocal microscope (Nikon TiE, Yokogawa CSU-X1) with temperature-controlled sample chamber (37C) and focus-locking mechanism for longer-duration recordings. Primary cultures were plated on 35mm MatTek dishes, and imaged in the same imaging buffer. We recorded for 15 minutes continuously at each field of view. The recordings were split into 3-minute segments due to memory constraints, and processed in a similar manner to the widefield recordings, as described below, with an additional step to remove periodic noise associated with spinning disk illumination (code and data supplement).

### Analysis of optical minis (Janelia)

Analysis of optical minis was automated using a constrained NMF framework with spatial components initialized from a correlation image (inspired by (43, 44)). Matlab code that performs these analyses is available in the code and data supplement. To perform factorization in a memory-efficient manner, each recording was first downsampled 2x in each spatial dimension and 8x in time by binning. F_0_ was then computed per-pixel and subtracted from the video to generate a downsampled baseline-subtracted video. A correlation image was computed where each pixel value is the mean correlation between that pixel and its 4-connected neighbors across time, after highpass filtering the video at 8 Hz. NMF was performed using a multiplicative updates algorithm, subject to constraints on the contiguity and sparsity of spatial components. The spatial factors were initialized as 2D gaussians (σ=530 nm) with a spatial extent of 2.8x2.8 µm, with no support outside this extent. A factor was added at each local maximum in the correlation image that exceeded a threshold value. An additional 20% of spatial factors were added to model background; these were not spatially constrained during optimization. Following factorization, spatial factors were merged if they were immediately adjacent or overlapping in space and had a Spearman rank correlation in time of at least 0.25. The spatial factors were then upsampled to the original spatial resolution and a final 20 rounds of multiplicative updates were performed on the full-resolution (space and time) data to generate a high-resolution factorization. From the resulting factorizations we computed the SNR and the detected event frequency of each site (see code). The SNR was defined as the amplitude of the third-largest peak in the highpass-filtered signal over the 1-minute duration of the recordings used for screening, divided by the standard deviation of noise estimated from the trace power spectrum. Detected event frequency was defined as the number of peaks exceeding 3 standard deviations of the noise. Detected site density is the number of detected sites with SNR>3 per unit area.

### Manipulations of synaptic release (Hoppa laboratory)

Hippocampal CA1-CA3 regions were dissected with dentate gyrus removed from P1 Sprague-Dawley rats of either sex (mixed litter), dissociated (bovine pancreas trypsin; 5 min at room temperature), and plated on polyornithine-coated coverslips (Carolina Biological; item 633095; 22x22x0.17 mm borosilicate glass) inside a 6 mm diameter cloning cylinder (Ace Glass) as previously described (45). Calcium phosphate transfection was performed on 5-day-old cultured neurons with plasmids encoding iGluSnFR variants, or for TeNT experiments, the same concentration of iGluSnFR-encoding plasmid plus a plasmid encoding TeNT (53). Treatment and control cultures (e.g. + and - TeNT) for each experiment were paired, i.e. used the same batches of neurons, plated and transfected at the same time, and cultured in the same incubator. Experiments were performed at 35° C using a custom-built objective heater. Coverslips were mounted in a rapid-switching, laminar-flow perfusion and stimulation chamber on the stage of a custom-built laser microscope. The total volume of the chamber was ∼ 150 μl and was perfused at a rate of 400 μl/min. During imaging, cells were continuously perfused in a standard saline solution containing the following in mM: 119 NaCl, 2.5 KCl, 2 CaCl2, 2 MgCl2, 25 HEPES, 30 glucose, solutions were supplemented with 10 μM 6-cyano-7-nitroquinoxaline-2,3-dione (Alomone) and 5-Phosphono-D-norvaline, D-Norvaline, 5-Phosphono-, D(-)-APV, D-2-Amino-5-phosphonovaleric acid, D-2-Amino-5-phosphopentanoic acid, D(-)-AP-5 (Alomone). For measuring exocytosis, neurons transfected with iGluSnFr variants were illuminated by a 488 nm laser 2 mW (Coherent OBIS laser) with ZET488/10x and ZT488rdc dichroic (Chroma) through a Zeiss EC Plan-Neofluar 40 x 1.3 NA Objective. iGluSnFr fluorescence emission was collected through an ET525/50m filter (Chroma) and captured with an IXON Ultra 897 EMCCD (Andor). iGluSnFr fluorescence was collected with an exposure time of 9.83 ms and images were acquired at 100 Hz. pHluorin-transfected cells were imaged under the same conditions as iGluSnFR variants, except with 8 mW of illumination intensity. Stimulation for firing action potentials for evoked vesicle fusion were evoked by passing 1 ms current pulses, yielding fields of ∼12 V/cm^2^ using platinum/iridium electrodes. Spontaneous release was easily identified by eye in absence of stimulation. These same sized signals can be identified in the presence of TTX (3 µM) confirming that they are spontaneous fusion (not shown). Images were analyzed in ImageJ (http://rsb.info.nih.gov/ij) by using custom-written plugins (http://rsb.info.nih.gov/ij/plugins/time-series.html). GluSnFR measurements were made from manually selected ROIs (2 micron diameter) drawn at individual responsive sites on isolated axons, then averaged across sites for each cell. This analysis of individual release sites differs from previous analyses of larger areas of mixed neurites (20). Individual cells were treated as the unit of variability; analysis of variance indivated no effect of culture batch on response amplitudes.

### Neuronal culture screen (Janelia)

Details on culture screening and analysis are provided in Supplemental Note 2. iGluSnFR variants were cloned into a mammalian expression vector and transfected into hippocampal and/or cortical primary cultures from neonatal (P0) pups. Imaging was performed in poly-D-lysine-coated 96-well glass-bottom plates. 14 days post transfection, culture medium was exchanged three times with 500 µL imaging buffer (145 mM NaCl, 2.5 mM KCl, 10 mM glucose, 10 mM HEPES, 2 mM CaCl_2_, 1 mM MgCl_2_, pH 7.3) and imaged in 75 µL imaging buffer and drug cocktail to inhibit synaptic transmission (10 µM CNQX, 10 µM (R)-CPP, 10 µM gabazine, 1 mM (S)-MCPG (Tocris)). Neurons were field stimulated with 1 and 20 pulses at 83 Hz, and imaged with 10X objective using an electron multiplying charge coupled device (EMCCD) camera (Hamamatsu ORCA Fusion C13440-20C, 1024x1024, center quad, 5.5 ms exposure, 182 Hz, 798 frames). Illumination was delivered by blue light (470/40 nm, Cairn Research Ltd; emission 525/50 using GFP filter cube). The approximate light density used was 0.34 mW/mm^2^ at 470 nm. Stimulation pulses were synchronized with the camera using data acquisition cards (National Instruments), controlled with Wavesurfer software (https://wavesurfer.janelia.org/). Imaging was performed at room temperature. Analysis of recordings was performed with custom Matlab code available in the software supplement (Supplemental Note 2). Time-integrated SNR (Figure 2e) was calculated by integrating the photon response over the duration that it exceeded the baseline (mean response over 80 samples prior to stimulus onset), and dividing by the shot noise associated with the baseline brightness over that period (i.e. the square root of the integrated baseline).

### 2P-excited bleaching measurements in vivo (Janelia)

To investigate photobleaching properties of new iGluSnFR variants *in vivo*, we imaged dendrites of excitatory cortical neurons in mouse visual cortex. Viral injections (100 nl, 250 µm below brain surface) were performed followed by implantation of a 4mm cranial window, as described previously (18). Imaging was performed on a customized adaptive optics two-photon microscope based upon a Thorlabs Bergamo II microscope. Excitation was provided from a Coherent Discovery NX TPC. The objective was a Olympus XLUMPFLN objective, 20x, 1.0 NA. A FOV of 133 x 67 µm was scanned at 156 Hz for the bleaching experiments. Emitted light was filtered through a 525/50 bandpass dichroic, collected via a ThorLabs PMT2100, and digitized via a laser-clocked National Instruments DAQ card controlled by ScanImage 2020. The imaging parameters were selected to match those we use in typical synaptic imaging experiments. Recordings of 12000 frames were collected at 156 fps with laser power of 33 mW at sample at a wavelength of 1010 nm, maximum depth of 100 µm below dura for both SF-Venus.iGluSnFR.A184V and iGluSnFR3.v857-PDGFR. Recordings were motion corrected (46) and pixels were selected that exceeded a threshold mean brightness over time. The traces for the selected pixels were then averaged to generate a single photobleaching trace. A constant background level, the mean recorded brightness in an unlabeled portion of the image, was subtracted for each field of view. A total of 34 FOVs (from 5 mice) for v857 and 27 FOVs (from 5 mice) for SF-Venus.iGluSnFR.A184V were analyzed.

### 1P in vivo imaging (Janelia)

We performed bulk imaging of iGluSnFR responses in the same mice transduced with AAVs encoding iGluSnFR variants as described for 2P-excited bleaching measurements, above. Four animals per construct were imaged at four time-points after surgery (3,4,5, and 6 weeks post-injection). In every animal there were four injection sites, two for a given iGluSnFR3 construct and two for SF-Venus.iGluSnFR.A184V serving as a within-animal control. A custom-build widefield microscope was used, incorporating an Orca Flash 4.0 Camera and a Thorlabs ITL200 tube lens. Each mouse received 4 non-overlapping AAV injections. Low-magnification fluorescence recordings of all injection sites were performed with an Olympus UPlanApo 4x objective, and sites were recorded individually through a 25x Olympus XLPlan N 1.05 NA objective. Excitation: SOLIS 470nm LED; hq 470/40 41109 Chroma Excitation filter. Et 560/40 319882 Chroma Emission filter. 512x512 16-bit images (4x pixel binning in both x and y) were acquired at 100 fps using Hamamatsu HC Image software. Illumination intensity was constant across all sessions. Drifting grating visual stimuli were presented using the same apparatus, grating spatial and temporal frequencies, and anesthesia conditions described in (18). Motion stimuli (2s duration) of 8 directions were presented in sequence spaced by 2s periods of luminance-matched gray screen, 8 times each.

Recordings were motion corrected (46) and background was subtracted by manually selecting a ROI within vasculature in each FOV. Pixels of interest were automatically detected based on their brightness and distance from automatically-detected vasculature (see code supplement). Pixels of interest were averaged to generate a single trace, separated into low-frequency (F0) and high-frequency (ΔF) components, and motion artefacts were then subtracted from the ΔF trace by regression of the low-pass filtered X and Y offsets obtained during motion correction (and corresponding quadratic terms) against the trace. Responses were averaged across stimulus presentations to generate a direction-independent mean response for each session. A subset of sessions showed very weak or no evoked modulation. We therefore sorted sessions for each site by power in the [0.2 0.3] Hz frequency interval and used only the top 50% of sessions per variant to calculate the grand mean response for each indicator.

Intrinsic signal imaging (ISI) was performed during the first and last imaging sessions for each animal to assess visual responsiveness at each recorded site over time. ISI was performed on the same microscope as fluorescence imaging, using a Mitutoyo 7.5x 0.21 NA objective, by collecting reflected light from a M625L4 LED (Thorlabs) illuminating the field of view from an angle outside the acceptance cone of the objective. This commonly-used geometry for ISI reduces collection of light reflected from glass surfaces. 16-bit full-field images were collected at 33.3 fps using Hamamatsu HC Image software. Light intensity was set to the maximum that avoided camera pixel saturation anywhere within the field of view. The same 8 stimuli used for fluorescence imaging were repeated over a period of approximately 30 minutes to generate ISI traces, which were then processed in a similar manner to fluorescence recordings.

### Mouse preparation (Kleinfeld laboratory)

Wild-type mice (C57BL/6J, males, >8 weeks old) were used for thalamocortical bouton imaging. Adult (>8 weeks old) transgenic mice (Scnn1a-Tg3-Cre, #009613, The Jackson Lab) were used for layer 4 dendritic spine imaging. Mice were anesthetized with isoflurane using a precision vaporizer, 3% (vol/vol) in oxygen for induction and 1–2% (vol/vol) for maintenance and given the analgesic buprenorphine subcutaneously (0.1 µg per gram body weight). Body temperature was maintained at 37 °C with a heating pad during anesthesia. The animal was placed in a stereotaxic frame. A 4-mm craniotomy was created over the right vS1 cortex (centroid at 1.5 mm posterior to the bregma and 3.4 mm lateral from the midline). Dura was left intact. A cranial window by a single 4-mm round coverslip (No. 1) was embedded in the craniotomy and sealed with cyanoacrylate glue (Loctite, cat. no. 401). Meta-bond (Parkell) was further applied around the edge to reinforce stability. A titanium head-bar was attached to the skull with Meta-bond and the remaining exposed bone was covered with dental acrylic (Lang Dental).

Virus injection was conducted before the craniotomy was made. To label the thalamocortical projections in vS1 cortex, Cre-dependent SF-Venus-iGluSnFR.A184V/ SF-iGluSnFR.A184V/ iGluSnFR3.v82.GPI/ iGluSnFR3.v857.GPI / iGluSnFR3.v857.PDGFR (AAV2/1.hSyn.FLEX.SF-Venus-iGluSnFR.A184V, original titer: 3.6E+13 GC/ml; AAV2/1.hSyn.FLEX.SF-iGluSnFR.A184V, original titer: 1.12E+13 GC/ml; AAV2/1.hSyn.FLEX.iGluSnFR3.v82.GPI, original titer: 8.46E+13 GC/ml; AAV2/1.hSyn.FLEX.iGluSnFR3.v857.GPI, original titer: 3.17E+13 GC/ml; AAV2/1.hSyn.FLEX.iGluSnFR3.v857.PDGFR, original titer: 2.56E+13 GC/ml ) and AAV2/1.hSyn.Cre virus (original titer: 3.1E+13 GC/ml) were diluted and mixed to the final titer of from 8.5E+12 to 1.1E+13 GC/ml for iGluSnFR virus and 3.4E+10 or 6.3E+9 GC/ml for Cre virus and then injected (50 nl, 10 nl/min) to the barreloids of ventral posteromedial nucleus in wild-type mice, 1.7 mm posterior to the bregma, 1.8 mm lateral from the midline and at a depth of 3.25 mm. To label the layer 4 dendrites and spines in vS1 cortex, virus of iGluSnFR3.v857.GPI (AAV2/1.hSyn.FLEX.iGluSnFR3.v857.GPI, original titer: 3.17E+13 GC/ml) was diluted to 1.06E+13 GC/ml and injected at a 45° angle into the vS1 of Scnn1a-cre mice to the target coordinates of 1.5 mm posterior to Bregma, 3.4 mm lateral from midline and 400 µm depth. For all injections glass pipettes (Drummond) were pulled and beveled to a tip at 30-µm outer diameter and a syringe pump (Kd Science, Legato 185) was used to control the infusion.

### In vivo AO2P imaging (Kleinfeld laboratory)

In vivo imaging was carried out after 4 weeks of expression and 3 days of habituation for head fixation. All imaging experiments used head-fixed awake mice under adaptive optics two-photon microscopy. Mice were anesthetized with isoflurane shortly and given a retro-orbital intravenous injection of 20 µl Cy5.5-dextran PBS solution to label the lumen blood vessels 30 min before imaging. AO correction was applied at the imaging depth below 350 µm using the method described previously (33). Briefly, direct wavefront sensing was performed through the labeled microvessels within a square region of 50 µm to 100 µm on edge, with the exact central location as the functional imaging ROI. The excitation wavelength for wavefront sensing was 1250 nm. The measured wavefront error was then applied to the deformable mirror during the functional imaging of the same ROI. Wavefront measurement and correction were repeated when switching to a different ROI. For functional imaging, the laser was tuned to 950 nm for SF-iGluSnFR.A184V, 1030 nm for SF-Venus-iGluSnFR.A184V and iGluSnFR3.v82.GPI and 1000 nm for iGluSnFR3.v857.GPI/PDGFR. Post-objective power was under 100 mW for all measurements.

### Vibrissa tracking and stimulation (Kleinfeld laboratory)

The vibrissae of the mice were trimmed 3 days before functional experiments, leaving only one vibrissa, that was C1, C2 or D1, whose corresponding cortical column had an optimal expression of iGluSnFR. The mice were head fixed to the imaging rig with a running disk. The experiments were carried out in the absence of visible light and the running disk was illuminated by two infrared LEDs (Thorlabs, M940L3) to generate a bright-field contrast for vibrissae tracking. A high-speed camera (Basler, acA1300-200um) was used to track the vibrissae at the frame rate of 500 Hz. Air-puff deflection was used for vibrissa stimulation. Pulse-controlled compressed air, 20-ms pulse width, 5 p.s.i. at the source, was delivered through a fine tube, which was placed parallel to the side of the mouse snout and 20 mm away from the targeted vibrissa. The frequency of the air puffs was from 2 Hz to 30 Hz. A motorized moving pole was used for dynamic pole touch experiments. The pole moves back and forth within a 5-mm range along the azimuthal direction at an average speed of 1.25mm/s. The positions of the vibrissa and pole were extracted using DeepLabCut (47) and custom code written in MATLAB.

### Single-cell electroporation and 2P imaging in mouse V1 in vivo (Konnerth laboratory)

For in vivo experiments, 50-55 days-old C57Bl/6 mice were implanted with a headplate for subsequent 2P glutamate imaging. Anesthesia was induced with 2% isoflurane in pure oxygen and maintained at 1.5% during surgery. The body temperature was continuously monitored and maintained by a heating plate at 37.5 °C. Both eyes were covered with ophthalmic ointment. After injecting a local anesthetic (2% xylocaine) and an analgesic (Metamizole, 200 mg/kg), a headplate was attached to the skull above the left hemisphere of the brain with dental cement.

The procedure for single-cell electroporation of plasmids was adopted from a previously described method with some modifications (48). The plasmid of either SF-iGluSnFR.A184S or iGluSnFR3.v857 was dissolved in an artificial intracellular solution (135 mM K-gluconate, 4 mM KCl, 10 mM HEPES, 4 mM Mg-ATP, 0.3 mM Na2-GTP, 10 mM Na-Phosphocreatine) at a final concentration of ∼100 ng/ul. A fluorescent dye (100 μM OGB-1) was also included to visualize the patch-pipette and to verify the success of electroporation. For electroporation, a 3 mm round craniotomy was made above the left primary visual cortex (V1), and a coverslip of a similar size was fixed onto the craniotomy with vetbond. The glass coverslip had a small perforation that allowed the access of a patch-pipette to the cortical tissue underneath. After electroporation, the perforated coverslip was replaced by an intact one and the edge of the craniotomy was completely sealed with Vetbond (3M).

Dendritic imaging of labeled neurons was performed with a custom-built 2P microscope. The microscope was equipped with a 12-kHz resonant scanner (Cambridge Technology) and controlled by LabVIEW. Excitation laser was a mode-locked Ti:sapphire laser (Mai Tai DeepSee, Spectra-Physics) and the average power below the objective (40x, NA 0.8, Nikon) was between 20-30 mW. Neurons expressing SF-iGluSnFR.A184S were imaged at 100 Hz frame rate with 940 nm excitation light and neurons labeled with iGluSnFR3 were imaged at 200 Hz with 950 nm excitation light. For visual stimulation, a 10.5-inch Samsung tablet was placed at a distance of 10 cm in front of the right eye of the mouse. The screen covered a visual space of 96 degree in azimuth and 70 degrees in elevation and had a mean brightness of 4.8 cd/m^2^. Full-field square wave drifting gratings (0.03 cycle per degree spatial frequency, 2 Hz temporal frequency, 2 s duration) were presented at 8 directions (45° step, in a fixed order) using a custom Android app.

For simultaneous 2P axonal imaging and cell-attached recordings, a z-stack of images of the labeled neuron was acquired and the axon was manually traced to identify unambiguously boutons from the same neuron. Before recording, a perfusion chamber was fixed on top of the headplate, and the coverslip on the cortex was changed to one with an opening. Warm artificial cerebro-spinal fluid (ACSF) (125 mM NaCl, 26 mM NaHCO3, 4.5 mM KCl, 2 mM CaCl2, 1.25 mM NaH2PO4, 1 mM MgCl2 and 20 mM glucose) was perfused throughout the experiment to keep the brain temperature constant. A patch-pipette filled with ACSF containing 25 μM Alexa Fluor 594 was approached under visual control to the soma of the labeled neuron and, once the tip of the patch-pipette touched the membrane, gentle negative pressure was applied to obtain a seal resistance of 40 MΩ. Once a stable cell-attached recording was established, spontaneous activity was recorded, while 2P glutamate imaging of axonal boutons was performed at a framerate of 500 Hz. Electrophysiological signals were acquired either in the current-clamp or in the voltage-clamp mode by a patch-clamp amplifier (EPC10, HEKA).

### Data analysis (Konnerth laboratory)

All data were processed and analyzed using MATLAB. Drifts of images in the xy-plane were corrected using a cross-correlation based registration algorithm. Regions of interest (ROIs) were automatically segmented based on the correlation image for each recording. The mean fluorescence signal F from each ROI was extracted and the fluorescence change was calculated as ΔF/F0 = (F-F0)/F0. For dendritic recordings with visual stimulation, F0 was defined as the mean of fluorescence signals during a period of two seconds before visual stimulation. For spontaneous activity recorded from axonal boutons, F0 was defined as the mean of fluorescence signals during all ‘No AP periods’. ‘No AP periods’ were the intervals between APs, starting 200 ms after the last AP. All traces displayed in Figure 5 were filtered using a Savitzky-Golay filter.

For the analysis of kinetics and specificity (Figure 5e), the traces from axonal boutons were denoised with a non-negative deconvolution algorithm (49) using the estimated range of tau for each sensor. ‘Single APs’ were APs that were separated by more than 200 ms from the preceding and following APs. The amplitude of a ‘single AP event’ was the peak fluorescence value during a period of 20ms following the electrically recorded spike. ‘No AP events’ were local maxima of the deconvolved trace during ‘No AP periods’.

Pixelwise orientation maps (Figure 5i, j) were generated as previously reported (18). Briefly, responses (mean fluorescence change) to 8 visual stimulation directions were obtained for each pixel. The responses were mapped onto orientation vector space and summed. The preferred orientation was the angle of the resultant vector. Orientation selectivity (OSI) was calculated as the length of the resultant vector divided by the sum of responses to each orientation. The response magnitude was defined as the 2-norm of responses to all directions. The hue of the orientation map encodes preferred orientation, the saturation corresponds to OSI, and the lightness is set by response magnitude and is normalized to the pixel with the largest magnitude from the map of v857.

### Expansion Microscopy (Janelia)

Female EMX1-cre mice 8 weeks old were injected in cortex at multiple sites in a 3.5mm craniotomy over V1 with 40nL of 5e11 GC/mL AAV2/1 encoding hSyn.Flex-iGluSnFR.v857 in the three backbones (PDGFR, GPI, and SGZ; 1 construct per mouse; 2 mice per construct). A cranial window was affixed to seal the craniotomy and allow expression to be confirmed before histology. Mice were transcardially perfused 16 days after virus injection, and gels prepared from labeled cortical regions using a protease-free variant of Ten-fold Robust Expansion (TREx) microscopy (32). Gels were labeled with primary and secondary antibodies as follows: Primary: mouse anti-bassoon #ab82958 (abcam), rabbit anti-homer #ab97593 (abcam), chicken anti-GFP # ab13970 (abcam). Secondary: goat anti-mouse AF594 A-11012 (Invitrogen), goat anti-rabbit Atto 647N 40839-1ML-F (Sigma Aldrich), goat anti-chicken AF 488 A-11039. For imaging, gels were expanded in purified water (Millipore Milli-Q) for 1 hour, placed in a glass-bottomed dish (coated in poly-L-lysine to prevent sample drift), and imaged using a Zeiss LSM800 confocal microscope using a 40x, 1.1NA water immersion objective. All images were acquired with the same image settings, except PMT voltages were adjusted to prevent digitizer saturation and adjust dynamic range across samples. Synapses were annotated manually by FT, and quality of annotations was verified blind to genotype by KP. The annotator was asked to: “*Draw a line connecting the presynapse to the post-synapse on an axis locally perpendicular to the iGluSnFR-labeled membrane, in the 2D image plane corresponding to the center of intensity of the pre- and post-synaptic labels*. *Only synapses with their axis in the plane of the image should be labeled.”* Annotations, image data, and software to visualize them (requires Matlab) are available in the data supplement.

### Statistics and reproducibility

Values and errors reported through the text (“### ± ###”) are mean ± SEM unless otherwise noted. Statistical analyses were performed in Graphpad Prism 8 (in vitro photophysics), and Matlab (others), as described in the text. Data and code used to produce the figures, and instructions for generating them, are provided in the data supplement.

### Reagent and availability

A variety of mammalian and bacterial expression plasmids for iGluSnFR3 variants v82 and v857 are available through Addgene (https://www.addgene.org/browse/article/28220233/). Requests for AAVs and other reagents can be made by contacting KP.

### Data availability

The data and code used to produce the figures are included in the code and data supplement. Requests for data can be made by contacting KP.

## Funding Support

All research performed at the Janelia research campus was funded by the Howard Hughes Medical Institute.

AJR, SJB, MBH: NSF CAREER 1750199, NIH R01NS112365, William H. Neukom Institute for Computational Science at Dartmouth College

RL, PY, XJ, DK: NIH U19 NS123717 and U24 EB028942

YC, MK, AK: The work was supported by the Deutsche Forschungsgemeinschaft (SFB 870) and the Max-Planck-School of Cognition. A.K. is a Hertie-Senior-Professor for Neuroscience.

## Acknowledgements

We thank Julia Kuhl for creating artwork depicting the 3-state model of iGluSnFR (Figure 1a, Supplemental Note 1) and C-terminal anchoring domains (Figure 4b). We thank Yusuke Nasu and Robert Campbell (UTokyo) for sharing plasmids containing various N-terminal secretion constructs (Supplemental Figure 9). We thank Tim Brown (Janelia), the Tool Translation Team (Janelia), and John Macklin (Janelia) for providing resources and access to instrumentation.

## Author Contributions

**Conceptualization:** Abhi Aggarwal, Yang Chen, Samuel J. Bergerson, Filip Tomaska, Manuel A Mohr, The GENIE Project Team, Loren Looger, Jonathan Marvin, Michael Hoppa, Arthur Konnerth, David Kleinfeld, Eric R. Schreiter, and Kaspar Podgorski.

**Data curation:** Abhi Aggarwal, Yang Chen, Filip Tomaska, Jeremy P. Hasseman, Daniel Reep, Getahun Tsegaye, Marinus Kloos, The GENIE Project Team, and Kaspar Podgorski.

**Formal analysis:** Abhi Aggarwal, Rui Liu, Yang Chen, Amelia J. Ralowicz, Samuel J. Bergerson, Filip Tomaska, Daniel Reep, Xiang Ji, Marinus Kloos, Michael Hoppa, Arthur Konnerth, David Kleinfeld, and Kaspar Podgorski.

**Funding acquisition:** The GENIE Project Team, Michael Hoppa, Arthur Konnerth, David Kleinfeld, Eric R. Schreiter, and Kaspar Podgorski.

**Investigation:** Abhi Aggarwal, Rui Liu, Yang Chen, Amelia J. Ralowicz, Samuel J. Bergerson, Filip Tomaska, Timothy L. Hanson, Jeremy P. Hasseman, Daniel Reep, Getahun Tsegaye, Pantong Yao, Marinus Kloos, Deepika Walpita, Ronak Patel, Manuel A Mohr, Boaz Mohar, The GENIE Project Team, and Kaspar Podgorski.

**Methodology:** Abhi Aggarwal, Rui Liu, Yang Chen, Amelia J. Ralowicz, Filip Tomaska, Timothy L. Hanson, Jeremy P. Hasseman, Daniel Reep, Getahun Tsegaye, Pantong Yao, Xiang Ji, Paul W. Tilberg, Boaz Mohar, The GENIE Project Team, Michael Hoppa, Arthur Konnerth, David Kleinfeld, Eric R. Schreiter, and Kaspar Podgorski.

**Project administration:** Abhi Aggarwal, Jeremy P. Hasseman, Michael Hoppa, Arthur Konnerth, David Kleinfeld, and Kaspar Podgorski.

**Resources:** Jeremy P. Hasseman, Daniel Reep, Getahun Tsegaye, Ronak Patel, Manuel A Mohr, The GENIE Project Team, Jonathan Marvin, Michael Hoppa, Arthur Konnerth, David Kleinfeld, Eric R. Schreiter, and Kaspar Podgorski.

**Software:** Yang Chen, Filip Tomaska, Timothy L. Hanson, Jeremy P. Hasseman, Daniel Reep, Getahun Tsegaye, Xiang Ji, The GENIE Project Team, and Kaspar Podgorski.

**Supervision:** Abhi Aggarwal, Jeremy P. Hasseman, Paul W. Tilberg, Michael Hoppa, Arthur Konnerth, David Kleinfeld, Eric

R. Schreiter, and Kaspar Podgorski.

**Validation:** Abhi Aggarwal, Jeremy P. Hasseman, Daniel Reep, and Kaspar Podgorski.

**Visualization:** Abhi Aggarwal, Rui Liu, Yang Chen, Amelia J. Ralowicz, Samuel J. Bergerson, Jeremy P. Hasseman, Michael Hoppa, Arthur Konnerth, David Kleinfeld, and Kaspar Podgorski.

**Writing - original draft:** Abhi Aggarwal and Kaspar Podgorski.

**Writing - review & editing:** Abhi Aggarwal, Rui Liu, Yang Chen, Amelia J. Ralowicz, Samuel J. Bergerson, Loren Looger, Michael Hoppa, Arthur Konnerth, David Kleinfeld, and Kaspar Podgorski.

## Supplemental Notes

### Supplemental Note 1. Kinetic effects on the spatial specificity of neurotransmitter indicators

We set out to develop iGluSnFR variants that better distinguish synaptic versus extra-synaptic glutamate signals. To better understand how kinetic properties of the indicator impact these signals, here we consider a simple reaction-diffusion system in which glutamate transients are released from point sources, and glutamate subsequently interacts with the expressed indicator. Consistent with our data and previous work (e.g., (50)), we model the indicator as having three states, described below. We model the extracellular interface as a 2D surface.

We define the *spatial specificity* of a signal as the ratio of a signal near the release site (at distance r = 50 nm) to that farther away (r = 1 um). We first consider the spatial specificity of the glutamate concentration itself. For a non-cooperative indicator, the spatial specificity of the ligand (i.e., glutamate) is an upper bound on the spatial specificity of the indicator (i.e., iGluSnFR). This is because a non-cooperative indicator has a linear or sublinear response to its ligand, meaning that a proportional increase in ligand results in at most a proportional increase in indicator signal. Since the ligand concentration close to the release site is always larger than that further away, the ratio of the indicator signal near the release site to that farther away is at most equal to the ratio of the concentrations (Figure SN1-1a).

**Figure SN1-1.**
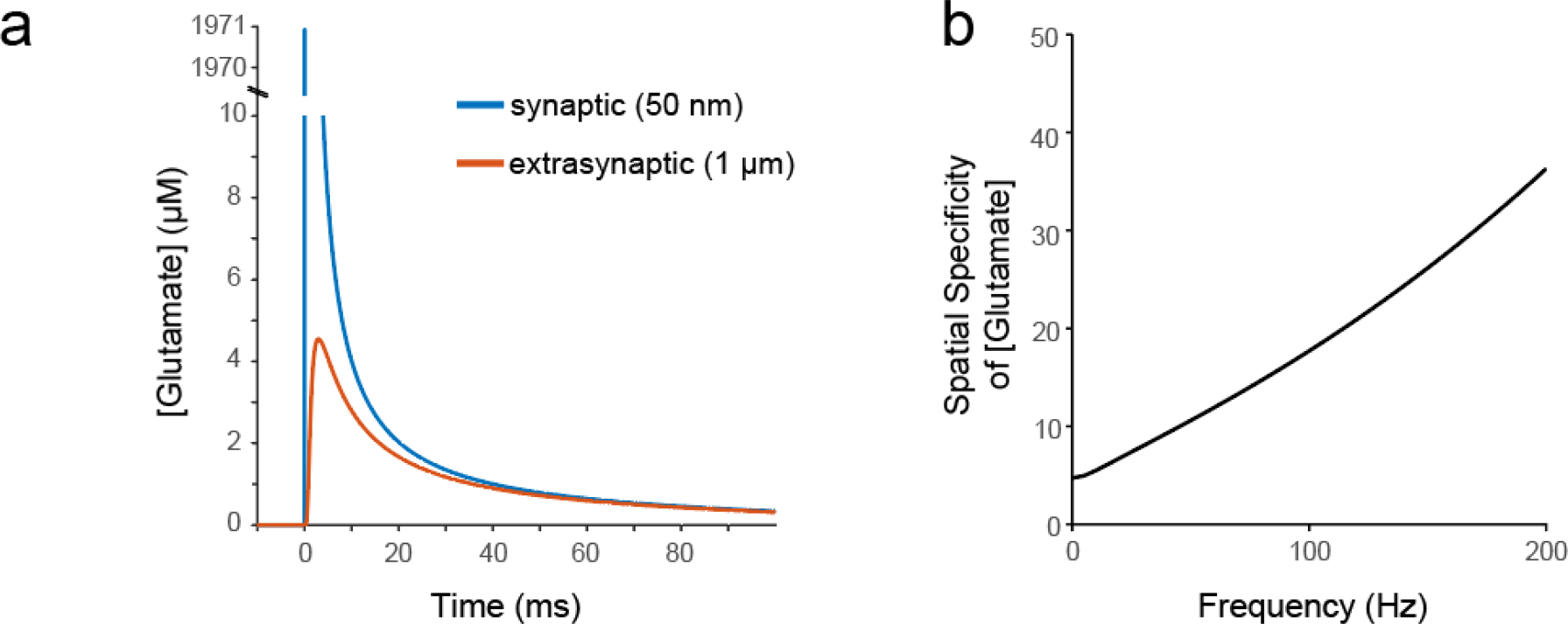
Simulations and measurements of iGluSnFR and glutamate kinetics. a) Simulated glutamate concentration at the synapse (50 nm from release site) and outside the synapse (1 µm from release site) in the absence of buffering. b) The ratio of the amplitude spectra of the signals in (a). This is the spatial specificity of glutamate as a function of temporal frequency. The spatial specificity of glutamate is an upper bound on the spatial specificity of any non-cooperative glutamate indicator.

The spatial specificity of the glutamate concentration can be characterized in the frequency domain (Figure SN1-1b). Faster components of the time-varying glutamate concentration are more specific for synapses than slower components. To better reproduce features of the glutamate signal that distinguish synaptic from extrasynaptic transients, the indicator should have fast rise and decay times, the imaging rate should be fast enough to capture those components, and data analysis should appropriately filter for higher-frequency components of the signal. At low frequencies, all indicators will show relatively low spatial specificity, because the glutamate concentration itself has low spatial specificity at those frequencies. This reasoning motivated the use of ΔF_fast_ as a screening criterion in neuronal culture (Supplemental Note 2).

iGluSnFR and its variants exhibit saturating ON rates, in which the formation of fluorescent species cannot exceed a maximum rate v_max_, presumably due to a rate-limiting conformational change (22, 50). ON rate saturation is characteristic of a kinetic model with at least 3 states, in which ligand binding (k_0-1_) rapidly forms a bound but dim species (S1), followed by a slower transition (k_1-2_) to a bright state (S2). This three-state model exhibits two different kinds of saturation phenomena. ON rate saturation occurs when large concentrations of ligand are rapidly applied, and unbound indicator (S0) becomes depleted in favor of bound-but-dim indicator (S1). This is distinct from saturation of the fluorescent state (S2), which occurs on slower timescales and involves depletion of the S1 state.

This 3-state model is closely related to the Michaelis-Menten model for enzymes. Using terminology of that model, we can define the asymptotic maximum initial ON rate v_max_ and the rate-half-saturating ligand concentration *K_M_* (*K_M_* = (k_1-0_ + k_1-2_)/k_0-1_). ON rate saturation is severe whenever the instantaneous glutamate concentration exceeds *K_M_*. At locations and times where instantaneous [glutamate] is higher, an indicator with saturating ON rates will under-report it more. Because instantaneous glutamate concentrations are much greater within the synapse than outside it, this phenomenon reduces the spatial specificity of the indicator (Figure SN1-2d).

**Figure SN1-2.**
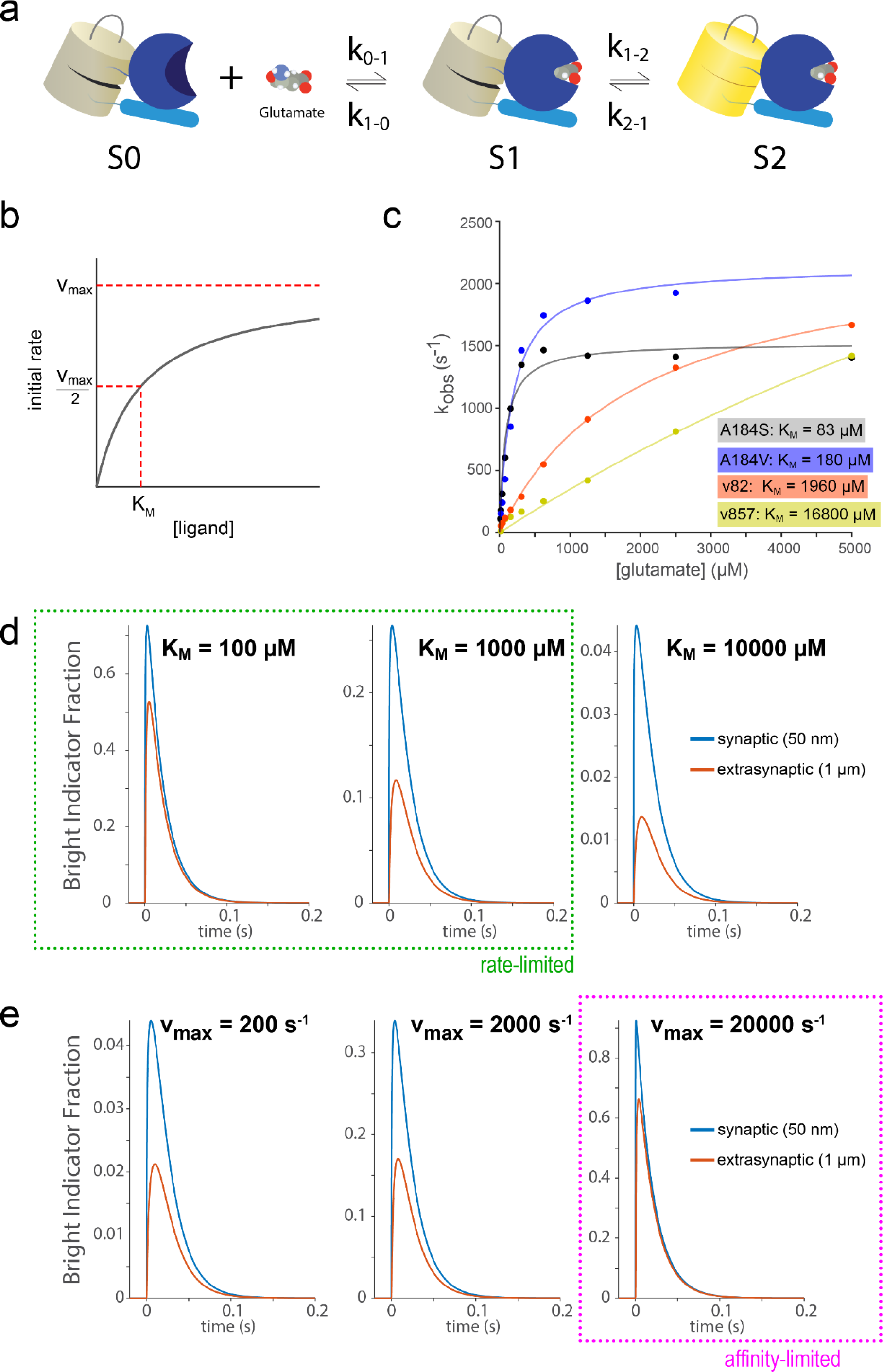
Modeling and measurements of iGluSnFR kinetics. a) Three-state model of iGluSnFR kinetics. In this model, initial glutamate binding is followed by a second, rate-limiting transition to the fluorescent state. b) Schematic of initial rate of formation of S2 in the three-state model as a function of ligand concentration. K_M_ is the rate-half-saturating ligand concentration. v_max_ is the asymptotic maximum ON rate. c) Measured initial rates (dots) for SF-iGluSnFR (A184V, A184S) and iGluSnFR3 variants at different glutamate concentrations and corresponding fits to the three-state model. Measurements were made in purified soluble protein at room temperature. d) Simulated fluorescence responses within and outside the synapse while varying *K_M_* and holding v_max_. constant (2500 s^-1^). e) Simulated fluorescence responses within and outside the synapse while varying v_max_ and holding *K_M_* constant (500 μM).

The effects of *K_M_* and v_max_ on spatial specificity are distinct . For a rate-limited indicator, increasing v_max_ (while holding *K_M_* fixed) does not improve its spatial specificity at low frequencies (Figure SN1-2e). However, increasing v_max_ results in faster indicator responses that can improve spatial specificity at high frequencies as described above (Figure SN1-1), and produces a higher bright state fraction for a brief impulse of glutamate (Figure SN1-2e).

Increasing v_max_ corresponds to increasing the association constant k_1-2_. As k_1-2_ increases relative to k_2-1_, the affinity of the indicator increases, and saturation of the bright state can occur in the model (‘affinity-limited’ indicators, magenta box in SN1-2e). This is a second form of saturation that in principle can reduce spatial specificity. However, experiments suggest that existing iGluSnFR variants are not affinity-limited: as long as initial binding is rapid (k_0-1_>>k_1-2_) an affinity-limited indicator must approach its maximum possible brightness during stimulation, but this does not occur for SF-iGluSnFR variants in culture as evidenced by field stimulation experiments followed by buffer exchange with varying concentrations of glutamate (20).

The above simulations suggest that the spatial specificity of existing iGluSnFR variants is limited by ON rate saturation, which is determined by kinetic parameters that vary substantially among existing variants (22, 50). These kinetic parameters are easily measured, and can therefore be engineered. On this basis, we set out to develop new iGluSnFR variants that are capable of reporting glutamate at high temporal frequencies, are not rate-limited (high *K_M_*), have large responses to brief glutamate impulses (high v_max_), and remain not affinity-limited.

### Simulation Methods

Simulations were performed using finite differencing methods in 2 dimensions, to model the geometry of the intercellular space. The MATLAB code for the simulations is available in the software package as `\`Figure 1 `and Supplemental Note 1\kinetics modeling\glutamateDiffusion.m`. The simulations reflect general principles of neurotransmitter sensing. Parameters used were selected to be relevant for glutamate sensing but are not meant to precisely model a particular indicator. Ligand reuptake and endogenous buffering are not modeled. The buffering effects of the indicator are modeled but not discussed here, for a discussion see (51).

## Supplemental Note 2. Directed evolution and multi-assay screen of iGluSnFR3

We began by subjecting WT (Round 0; **Figure SN2-1a**) to error prone polymerase chain reactions (EP-PCR; mean rate 4.5 to 9 DNA substitutions/clone) to generate mutant libraries of the soluble iGluSnFR coding sequence that were then assembled into a pRSET plasmid vector (Step 1). These libraries were transformed into electrocompetent *E. coli*, and plated onto agar plates containing ampicillin. In each round, we screened roughly 50 agar plates (∼10^5^ colonies) using either 2-photon (1030 nm) or 1-photon excitation (488 nm), and selected the brightest colonies by eye (Step 2) through laser-safety glasses. Macroscopic 2-photon excitation (∼2x4 cm) was achieved using a low repetition rate, high-power femtosecond fiber laser (1 MHz repetition rate, 55 W average power, 120 fs pulsewidth; Tangerine HP2, Amplitude Systemes) that was shaped into a line focus and scanned along one dimension at 500 Hz using a galvanometer mirror scanner (**Figure SN2-1b**). The selected bacterial colonies were inoculated into Studier’s autoinduction medium, incubated overnight, and washed with repeated cycles of centrifugation, decanting, and resuspension in glutamate-free PBS to remove glutamate. Bacteria were lysed using freeze-thaw cycles of liquid nitrogen (-196 ℃) and warm water bath (40 ℃) (see methods) (Step 3). Bacterial lysis by other means (i.e., chemically using B-PER, mechanical disruption, or sonication) resulted in lower protein yield, higher assay variability, and/or reduced ΔF/F_0_, as assessed using repeated measurements of the WT indicator (data not shown).

**Figure SN2-1.**
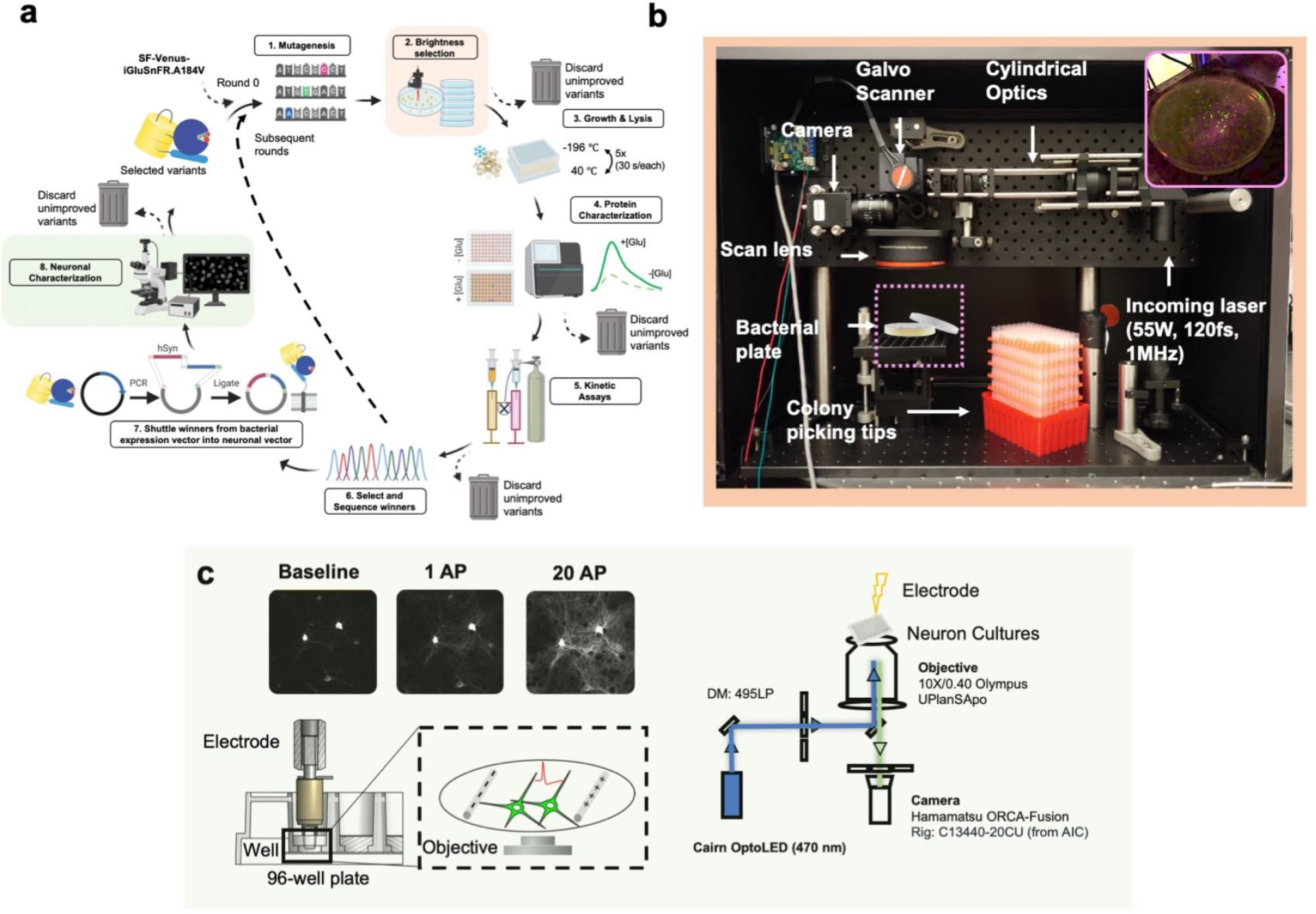
a) Schematic of multi-assay directed evolution screen. The smaller loop (steps 1-6) was performed for 20 generations, followed by the larger loop (steps 1-8) for an additional 2 generations. b) Custom large-area two-photon excitation system used to select colonies for 2P-excited brightness. Inset shows an example plate with fluorescent colonies excited by the system. The infrared excitation beam appears as magenta in this photo, but is invisible to the human eye. c) Schematic of in-neuron culture screening using the GENIE Project apparatus. Top, example frames showing a variant’s baseline and responses to 1 AP and 20 AP in the culture assay. Bottom, primary cultures were plated in a 96-well imaging plate, imaged using an inverted microscope and stimulated by a motorized dipping electrode (reprinted from (24)). Right, schematic of the culture assay imaging path. Schematic in (a) was created using BioRender.com (2021).

Clarified bacterial lysates were used for multiple assays (Step 4). In all rounds, a plate reader was used to measure fluorescence excitation spectra before and after the addition of glutamate. In most rounds, glutamate was added to a final concentration of 500 uM, but after the E23D mutation became fixed in our variant pool, we used 200 uM [glutamate] to select for higher-affinity variants. In lysate assays, variants were selected on the basis of the combination of ΔF/F_0_, excitation spectrum (we selected for variants with red-shifted peak excitation relative to EGFP), and glutamate-bound brightness in the plate reader assay (which may reflect intrinsic brightness of the indicator variants, but also properties such as maturation rate and bacterial copy number). DNA was then isolated and sequenced for the selected variants, and the rest were discarded (Step 6).

Following the first round of selection, the retained population of variants were recombined through staggered extension PCR (StEP) (52), then subjected to the same screening and selection process (Steps 2 to 6). Mutagenesis in subsequent rounds proceeded using diversification (via EP-PCR, site-saturation, site-directed mutagenesis, or combinatorial mutagenesis) and recombination, in alternating rounds.

From the third round onward, we performed additional assays in lysate and purified protein to further remove undesired variants from the pool. In lysate, we performed pH titrations in the presence and absence of glutamate, discarding variants with high pKa in the glutamate-bound state. In purified protein, we measured 2-photon excited molecular brightness at 1000 nm excitation, discarding variants with low brightness, and we measured in vitro kinetics in a stopped-flow assay to select for variants with high v_max_ and non-saturating k_obs_ (indicative of high K_M_) (Step 5). Additionally, we performed k_d_ titrations, and evaluated extinction coefficient, quantum yield, and molecular brightness in glutamate-free and glutamate-bound states. In each case, the sequential selection we performed ensured that all retained variants jointly met all selection criteria.

After 20 rounds of in vitro selection, a library of 120 variants were subcloned into a neuronal surface display vector (Step 7). These were then characterized in cultured rat neurons (Step 8) using a modified form of the established GENIE Project pipeline for high-speed imaging of field-stimulated neuron cultures in the presence of post-synaptic receptor antagonists to control the number and timing of evoked action potentials (24). This first neuronal screen identified v82 and several related variants as high-sensitivity indicators with similar on-neuron kinetics to WT. The soluble forms of these variants were further diversified and screened in bacteria and lysate, for brightness, response amplitude, and spectral properties. 812 variants were selected from this pool, generated as membrane-tagged mammalian expression plasmids, and screened in a second round of neuronal culture screening. After the second round of screening, the assay was repeated for selected variants with additional stimulus conditions (1,5,10,20,160 APs; Figure 2 of main text).

For neuronal culture screening, the GENIE Project apparatus was modified to use a high-speed camera (180fps, Hamamatsu ORCA Fusion) and to automatically focus on labeled neurites at the surface of the coverslip for each well. Each variant was transfected 4 times and plated into 16 wells. Each well was recorded while stimulating 1 and 20 AP field stimuli. We characterized a variety of properties for each well, including baseline brightness, photostability, and for both 1 AP and 20 AP conditions: the number of responsive pixels, the fractional change in brightness (ΔF/F_0_), SNR, and response onset and offset kinetics (Figure SN2-2). Summary data and code used for analysis is available in the data supplement.

**Figure SN2-2.**
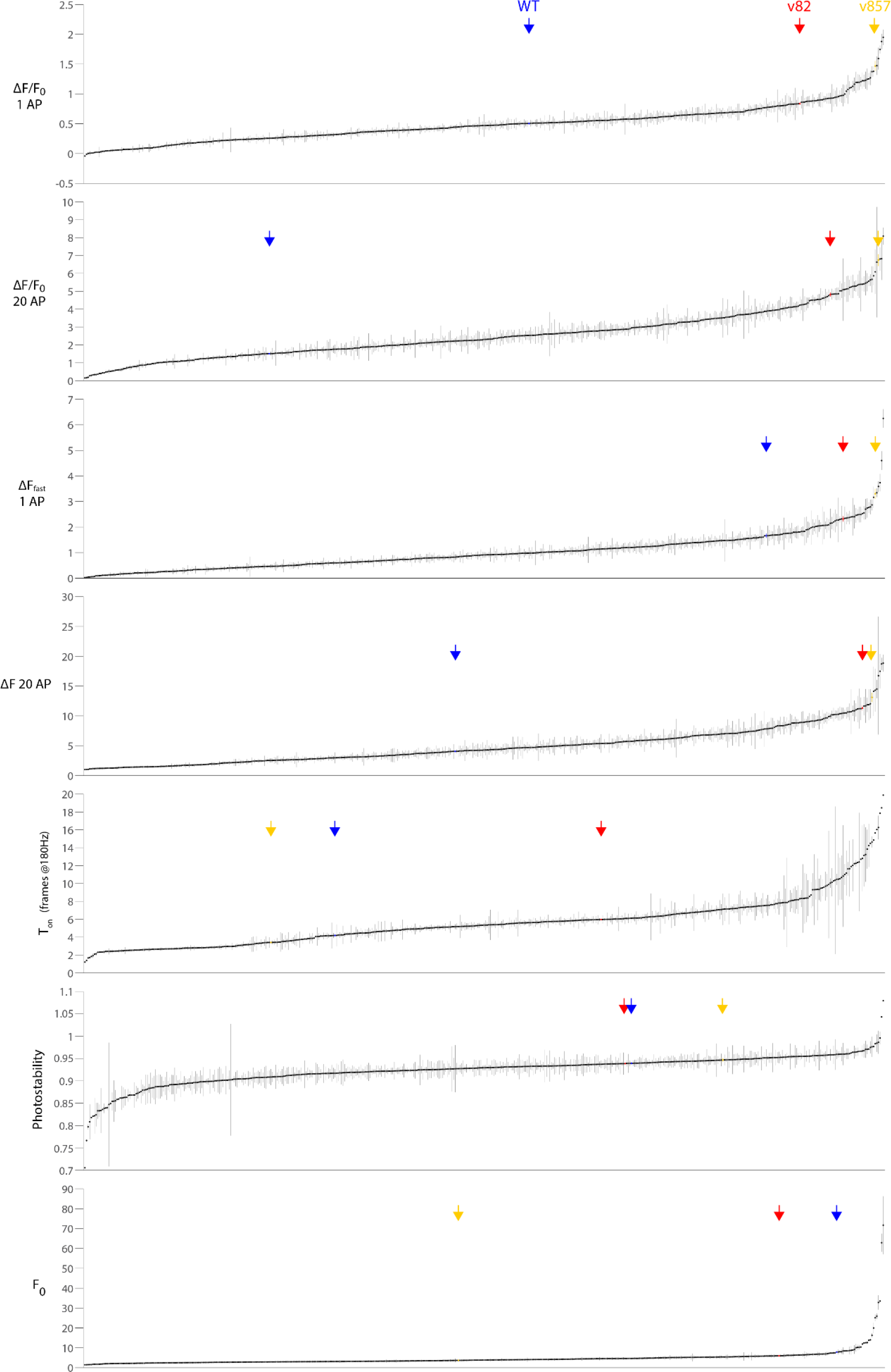
Properties of variants measured in the second round of neuron culture screen. Dots denote individual variants, sorted according to the plotted value. 499 variants are plotted; an additional 310 variants did not meet the threshold for minimum number of significantly responding pixels per well (300 labeled pixels exceeding a Z-score of 3.5). Error bars denote standard error of the mean across wells (N=14-16 wells per variant). Blue, Red, and Yellow arrowheads (and corresponding colored data points) denote values for WT, v82, and v857 variants, respectively. ΔF_fast_ (arbitrary units) is proportional to the excess number of photons collected in a 6-frame window following the stimulus in the top 10% of responding pixels. T_on_ is the calculated 10-90% rise time of the 1AP response, fit with sub-frame resolution using linear interpolation. v857 was selected in this screen on the basis of its desirable combination of properties. Photostability was calculated as the fraction of original intensity retained following 2 seconds of illumination; it is a measure of rapid photobleaching and/or photoswitching.

An important criterion we used to select variants in the cultured neuron assay was ΔF_fast_, the excess number of photons collected per pixel within 33 ms (6 frames) of 1AP stimulation, compared to a baseline before stimulation, averaged over the top 10% most responsive pixels in the same time window that met a minimum brightness threshold. These 10% most responsive pixels were selected using a separate stimulus presentation to avoid selection bias. The motivation for this criterion was to enrich measurements for synaptic glutamate signals. We reasoned that an indicator with increased spatial specificity at synapses should show large signals at a fraction of labeled pixels (those close to release sites), and would potentially show smaller signals at the remaining pixels. Moreover, this spatial difference should be most apparent over a short period of time following release (see Supplemental Note on Kinetics). Example responses for several variants are plotted in Figure SN2-3, for both the selected 10% of pixels (solid lines), and for all responding pixels (dashed lines). A large difference between the solid and dashed traces indicates that a subset of highly responsive pixels exhibit larger responses than the mean labeled pixel, which is expected for a more spatially specific indicator.

**Figure SN2-3.**
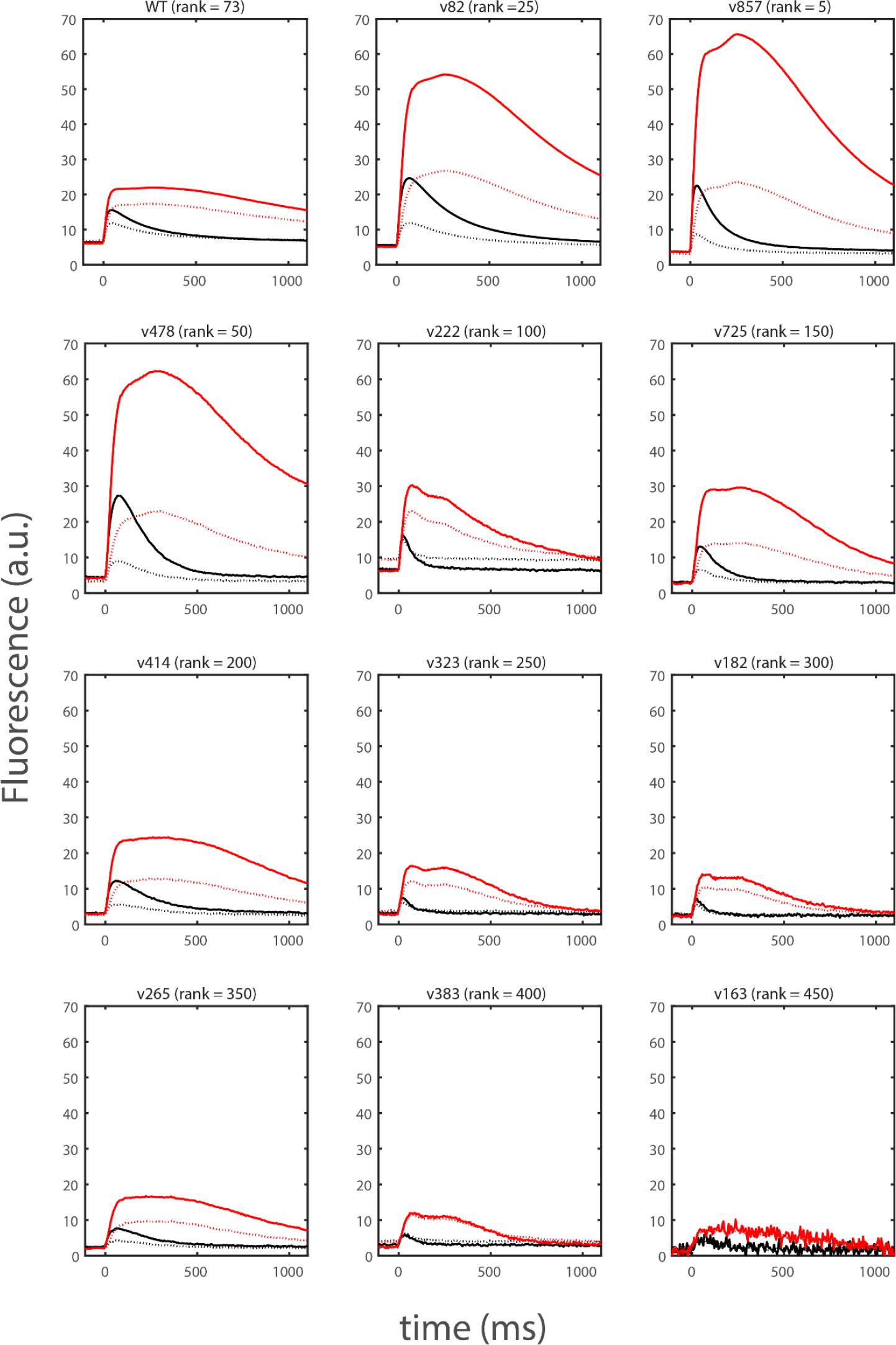
Example traces for variants measured in the second round of neuron culture screen. The black and red traces denote responses to 1 and 20 APs, respectively. Solid and dashed traces denote mean responses of the top 10% of pixels selected from each well (as described in text) and all labeled pixels, respectively. The rank of each variant in ΔF_fast_ within the screening round is displayed. The displayed traces show ∼1 second of the 5-second total recording duration.

## Supplemental Note 3. Mutations present in iGluSnFR3 variants

Variant v82 has 13 mutations relative to WT (with positions numbered from the opening of the iGluSnFR coding sequence^1^): E23D, N33S, Q35H, D43E, F82Y, V182S, N196D, S252N, N254I, R273H, A332T, I334N, T335S. Variant v857 has 15 mutations relative to WT (mutations relative to v82 are underlined): E23D, N33S, Q35H, D43E, F82Y, V182S, N196D, S252N, <u>N254T</u>, R273H, A332T, I334N, T335S, <u>L395F</u>, <u>G414S</u> (Figure SN3-1). The sites E23, V182, and G414 have been identified in previous studies (19,20,22), and the others are novel.

**Figure SN3-1.**
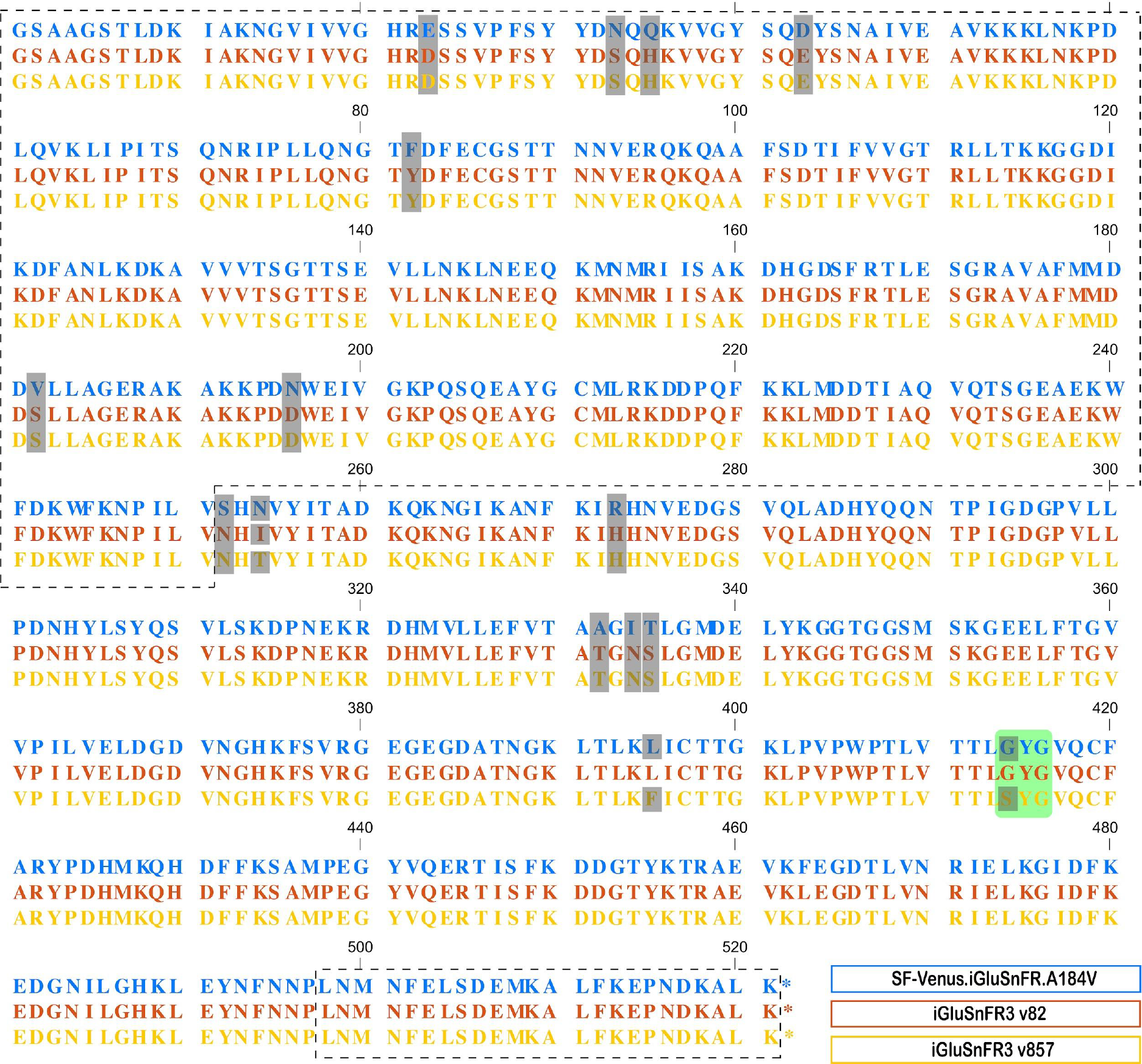
Sequence alignment of WT (blue), v82 (red), and v857 (yellow). Grey highlights denote mutations with respect to WT. Green box highlights the chromophore. Dotted outlines denote regions derived from gltI.

Notably, even though mutagenesis was performed across the entire coding sequence, most of the mutations selected by screening lie near a putative interface between iGluSnFR’s circularly-permuted fluorescent protein domain and the large (N-terminal) fragment of gltI (Figure SN3-2). No functionally-relevant point mutations were identified in the small (C-terminal) α-helix fragment of gltI. Truncation of this fragment in soluble iGluSnFR v857 (at residue 498) yielded a functional sensor, albeit with a diminished dynamic range (data not shown).

**Figure SN3-2.**
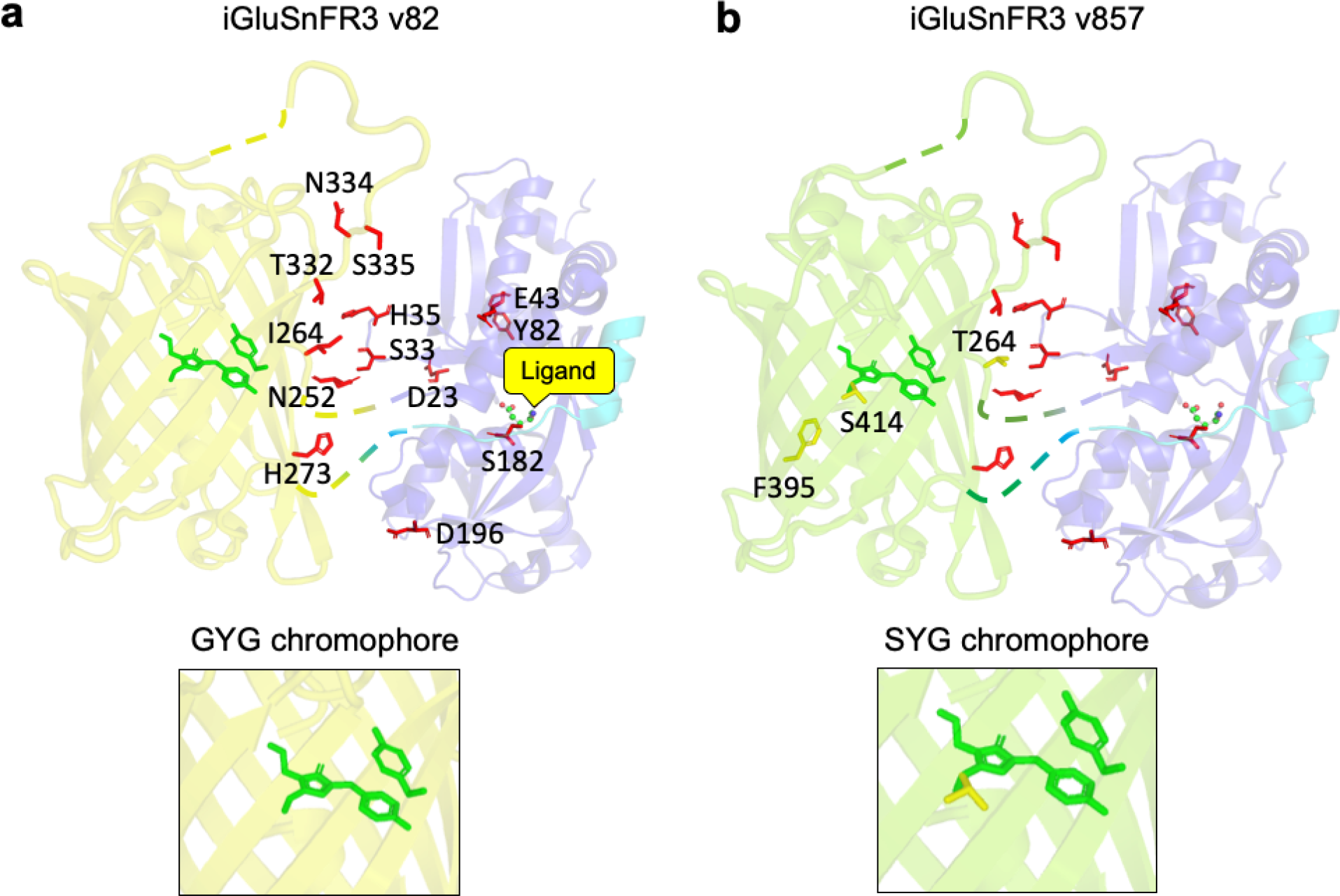
Crystal structures of YFP (PDB: 1F0B) and gltI (PDB: 2VHA) overlaid with mutations present in a) v82 and b) v857. Mutations are shown in red sticks. Zoom in boxes show the distinct chromophores of v82 and v857.

These mutations were identified over multiple rounds of screening involving multiple assays. Early rounds of screening relied exclusively on random mutagenesis and StEP recombination and resulted in a population of variants with high brightness, high ΔF/F_0_ and micromolar *K*_d_ containing the mutations E23V, N33S, Q35H, D43E, P75Q, F82Y, N196D, S252N, R273H, A332T, I334N, T335S, F463L. The selected population of variants at this point all exhibited rapidly-saturating, slow ON kinetics in purified protein (not shown). To diversify kinetic properties and enable selection for variants with faster and less-saturating ON kinetics, we then rationally mutated sites previously shown to affect iGluSnFR kinetics: E23 (1), and S70 (1, 2), on top of the selected population. We produced 40 variants mutated at these two sites, which we characterized in purified protein using a stopped-flow kinetics assay. In these variants, E23D improved the desired kinetic parameters, while E23V, E23A, and E23K significantly improved sensitivity but harmed ON kinetics (data not shown). Mutants of this pool at position S70 did not result in any fluorescent bacterial colonies. The E23D mutation reduced the affinity of the soluble sensor to an undesirable degree. We therefore further diversified this pool by adding the affinity-increasing mutation A182S (A184S in Marvin et al) to E23D-containing variants. In soluble protein, we found that addition of A182S to E23D variants increased their affinity approximately 10-fold, without significantly altering ON rates. The resulting population of variants formed the first pool (n=120 variants) screened in neuron culture. We found that variants containing both E23D and A182S mutations showed the largest rapid-onset (0-20 ms post-stimulus) responses to 1 AP field stimuli, our performance criterion in the neuron screen (Supplemental Note on In-Neuron Screening). The best of these was the variant v82. Kinetic characterization of v82 confirmed the variant’s improved K_M_ (Figure 1). Further diversification, consisting of saturation mutagenesis at sites selected for in earlier rounds of screening and error-prone PCR, resulted in identification of the additional mutations <u>N254T</u>, <u>L395F</u>, <u>G414S</u> present in v857. Kinetic characterization of v857 confirmed this variant’s even higher K_M_, in addition to other desirable properties identified in the multi-assay screen (Figures 1-3).

To assess functional relevance of the 15 mutations in v857, we individually reverted each single amino acid mutation to its WT identity. We evaluated each reversion in purified soluble protein, on the basis of excitation and emission spectra, *in vitro* ΔF/F_0_ to saturating glutamate, and affinity (Table 1). In all cases, reversion decreased *in vitro* ΔF/F_0_, or in case of S182V, resulted in no fluorescence in the purified protein. As expected, reversion of the chromophore mutation S414G resulted in a bathochromic shift, while the remaining positions had no effect on the excitation or emission maxima. These results suggested to us that all of the accumulated mutations in v857, when assessed independently and assayed *in vitro*, are necessary for optimal function of the sensor.

**Table SN3-1.** Measurements of all single-residue reversions of iGluSnFR v857 in soluble purified protein. NF = non-fluorescent in bacteria. Values at mean ± SEM. n = 3 for each measurement.

| Mutant | Exc<sub>max</sub> (nm) | Em<sub>max</sub> (nm) | $\Delta F/F_0$ | K<sub>d</sub> (μM) |
| --- | --- | --- | --- | --- |
| SF-Venus-iGluSnFR.A184V | 512 | 530 | 2.9 ± 0.1 | 24.4 |
| iGluSnFR3 v857 | 502 | 522 | 54.0 ± 2.6 | 195.9 |
| iGluSnFR3 v857 + D23E | 502 | 522 | 17.5 ± 0.3 | 70.9 |
| iGluSnFR3 v857 + S33N | 502 | 522 | 13.6 ± 0.6 | 307.3 |
| iGluSnFR3 v857 + H35Q | 502 | 522 | 39.5 ± 1.3 | 180.1 |
| iGluSnFR3 v857 + E43D | 502 | 522 | 10.7 ± 0.1 | 106.4 |
| iGluSnFR3 v857 + Y82F | 502 | 522 | 2.7 ± 0.2 | 60.0 |
| iGluSnFR3 v857 + S182V | NF | NF | NF | NF |
| iGluSnFR3 v857 + D196N | 502 | 522 | 5.5 ± 0.5 | 137.3 |
| iGluSnFR3 v857 + N252S | 502 | 522 | 10.6 ± 0.2 | 271.6 |
| iGluSnFR3 v857 + T254N | 502 | 522 | 11.4 ± 0.8 | 164.8 |
| iGluSnFR3 v857 + H273R | 502 | 522 | 28.6 ± 1.1 | 238.5 |
| iGluSnFR3 v857 + T332A | 502 | 522 | 4.2 ± 0.2 | 117.9 |
| iGluSnFR3 v857 + N334I | 502 | 522 | 18.1 ± 0.6 | 70.6 |
| iGluSnFR3 v857 + S335T | 502 | 522 | 13.6 ± 1.3 | 24.9 |
| iGluSnFR3 v857 + F395L | 502 | 522 | 15.9 ± 0.7 | 172.5 |
| iGluSnFR3 v857 + S414G | 510 | 524 | 7.0 ± 0.3 | 66.0 |

In addition, we have curated a database of mutations acquired throughout our multi-assay screen, with measurements of ΔF/F_0_, *K*_d,_ k_obs_, with the parameters measured in bacterial lysate or in purified protein. These measurements are available at the GitHub repository: https://tlh24.github.io/glusnfr-analysis/

## Supplemental Figures

**Supplemental Figure 1.**
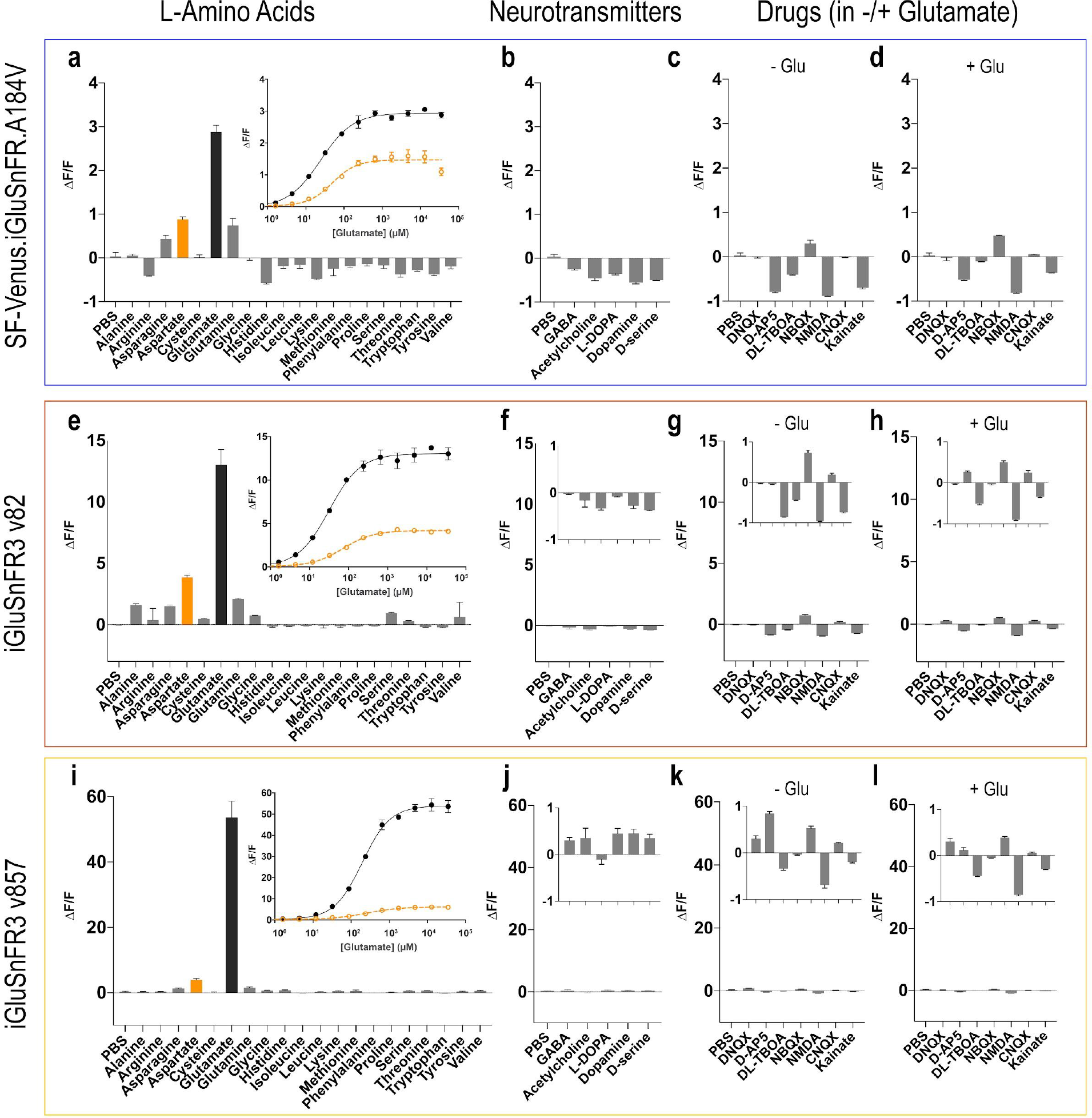
I*n vitro* specificity of soluble proteins SF-Venus-iGluSnFR-A184V (blue, top), iGluSnFR3.v82 (red, middle) and iGluSnFR3.v857 (yellow, bottom). a,e,i) ΔF/F_0_ of the three biosensors for additions of L-amino acids (10 mM concentration, pH 7.3, buffered in PBS). Zoom-in shows titrations for glutamate (black) and aspartate (orange). b, f, j) ΔF/F_0_ for additions of common neurotransmitters (GABA, Acetylcholine, L-DOPA, Dopamine, D-serine; 10 mM concentration, pH 7.3, buffered in PBS). c, g, k) ΔF/F_0_ for additions of glutamatergic drugs (DNQX [10 mM], D-AP5 [500 µM] , DL-TBOA [4 mM], NBQX [1 mM], NMDA [10 mM], CNQX [4 mM], Kainate [4 mM]) in the absence of glutamate and d, h, i) in presence of glutamate (10 mM, pH 7.3).

**Supplemental Figure 2.**
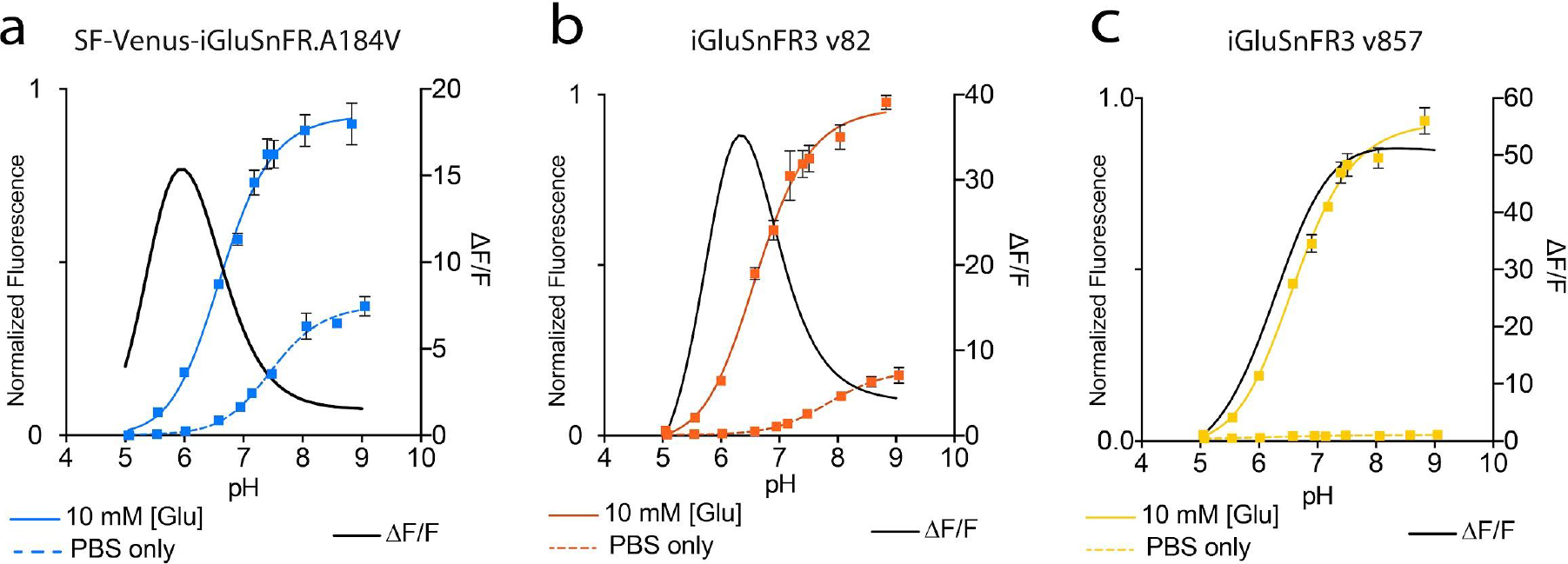
I*n vitro* pH titration of soluble proteins a) SF-Venus-iGluSnFR-A184V (left), b) iGluSnFR3 v82 (middle) and c) iGluSnFR v857 (right). Solid lines: saturating glutamate (10 mM, pH 7.3 buffered in PBS); Dotted lines: absence of glutamate. Sigmoidal fits are overlaid. Black lines show ΔF/F_0_ as a function of pH. All measurements were made using purified soluble protein; N=3 for each measurement, error bars represent SEM.

**Supplemental Figure 3.**
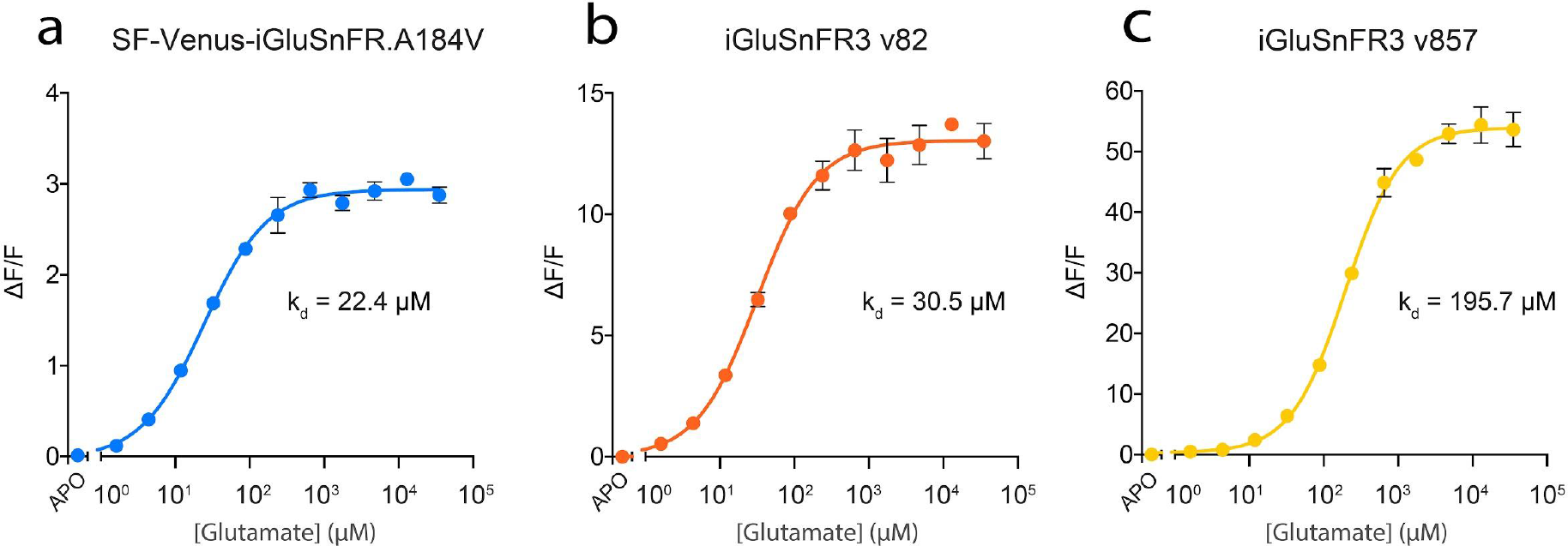
I*n vitro* glutamate titration of soluble proteins a) SF-Venus-iGluSnFR.A184V, b) v82 and c) v857 in soluble protein (pH 7.3), with corresponding fits and dissociation constants (K_d_). N=3 for each data point, error bars represent SEM.

**Supplemental Figure 4.**
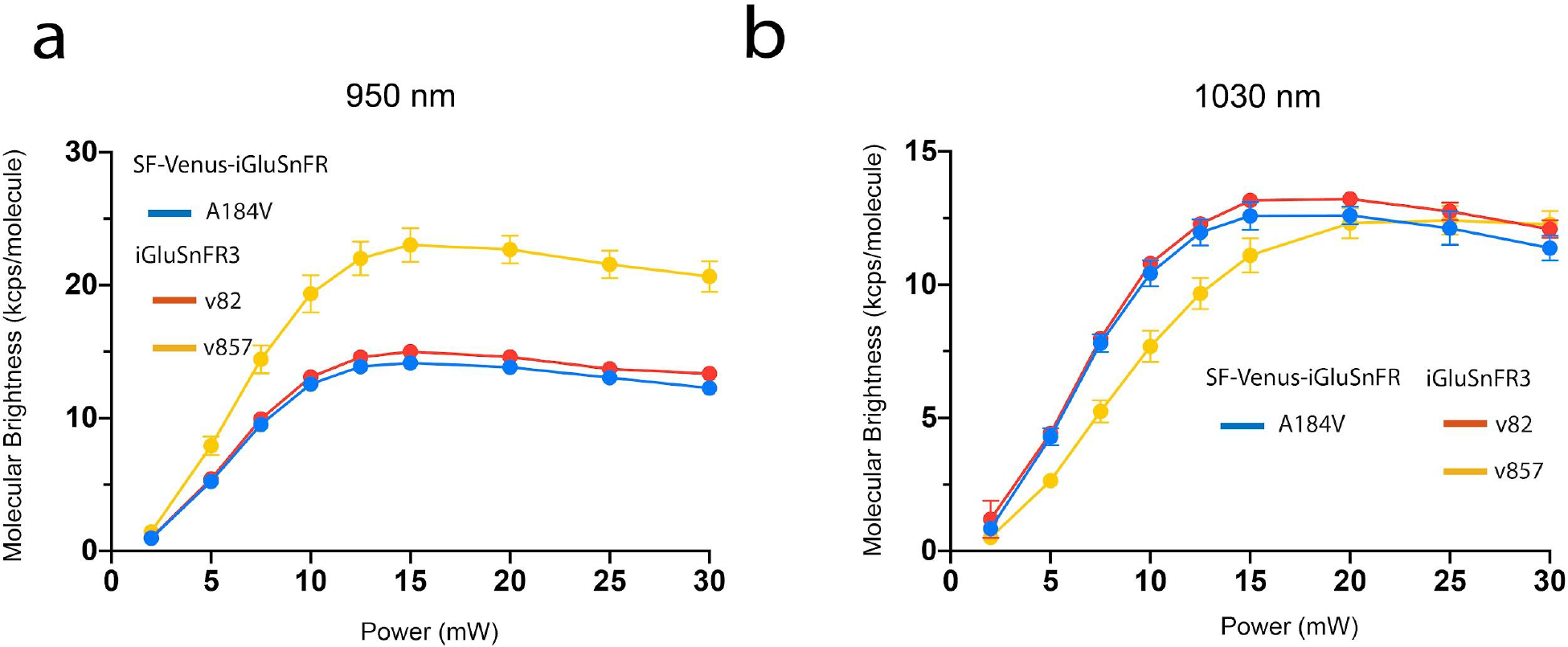
Molecular brightness (kcps/molecule) of iGluSnFR variants (blue - SF-Venus-iGluSnFR.A184V; red - iGluSnFR3.v82; yellow - iGluSnFR3.v857) measured using fluorescence correlation spectroscopy (FCS) at a) 950 nm and b) 1030 nm at varying power (mW). All measurements were made using purified soluble protein; N=3 for each measurement, error bars represent SEM.

**Supplemental Figure 5.**
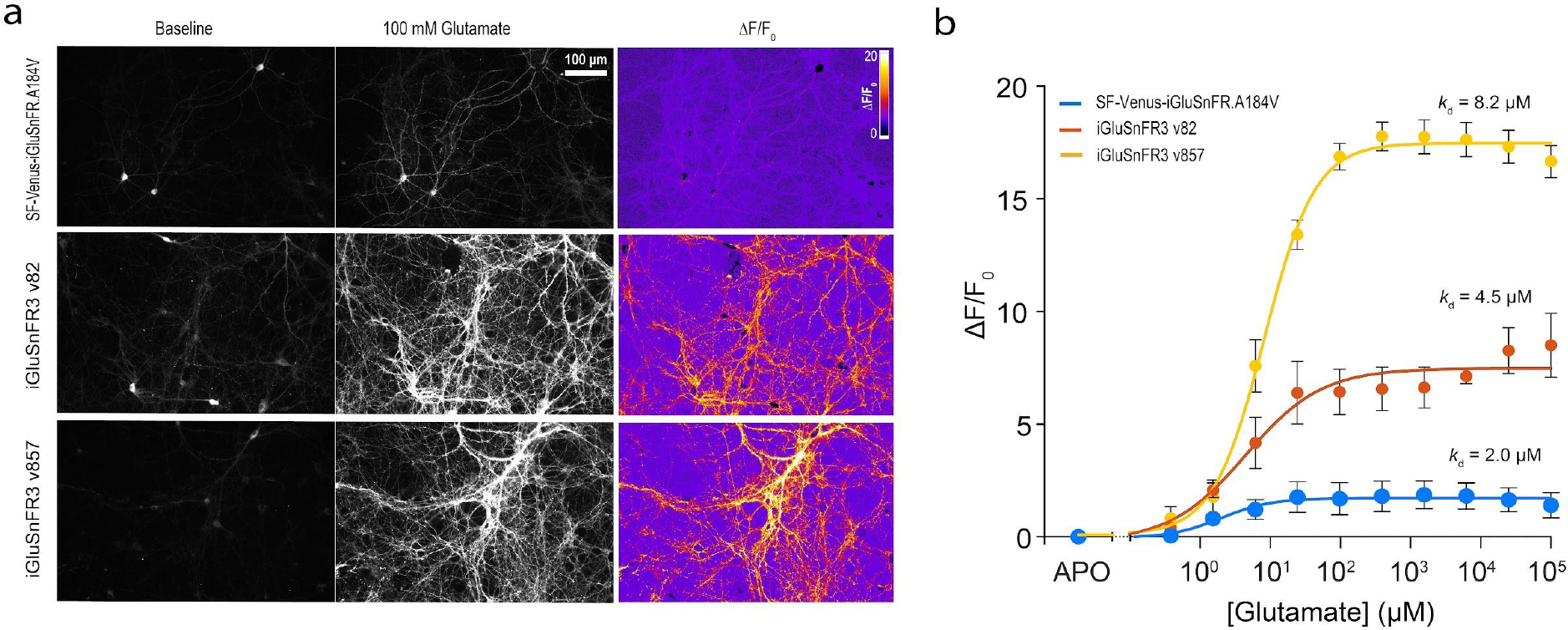
a) Glutamate titrations of variants SF-Venus-iGluSnFR-A184V, iGluSnFR3.v82 and iGluSnFR3.v857 on the surface of cultured hippocampal neurons. Baseline, 100 mM glutamate, and ΔF/F_0_ heat maps shown. b) ΔF/F_0_ affinity curves. ‘APO’ represents the baseline without any glutamate. ΔF/F_0_ is computed by using the average intensity of the entire field of view (FOV) and dividing by average intensity of APO, after background subtraction. The points are fit using a sigmoidal curve. Each data point is an average of N=3 FOVs from independent wells, and error bars represent SEM.

**Supplemental Figure 6.**
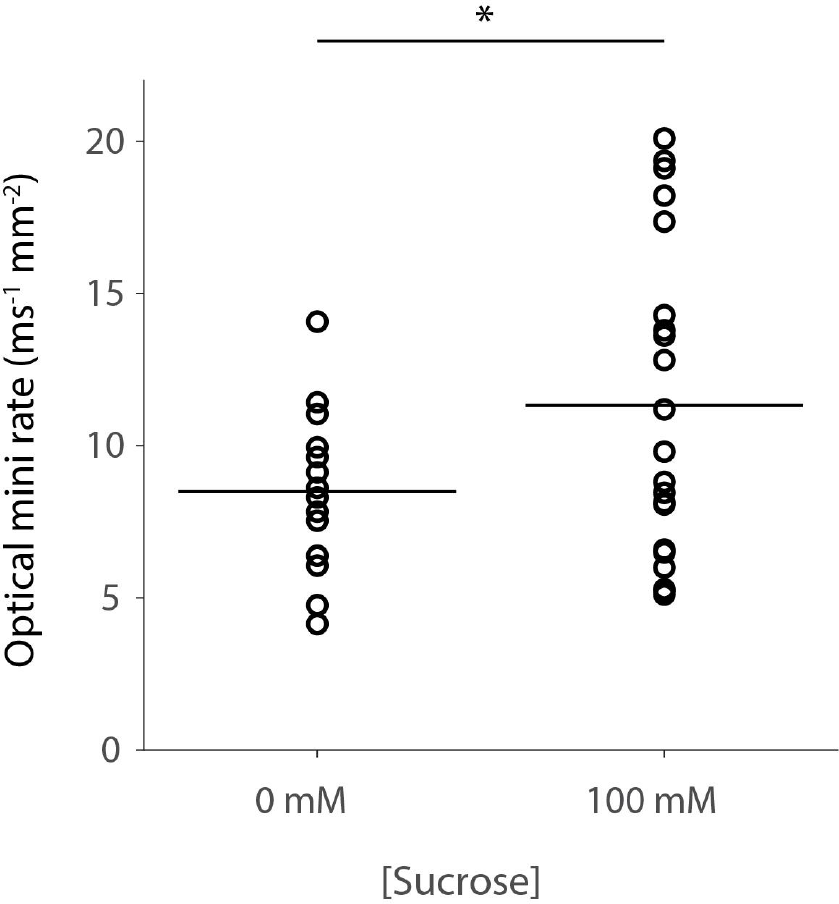
Effect of osmolarity on optical mini rate in primary cultures expressing iGluSnFR3.v857. Rate of detected optical minis in imaging buffer before (left) and after (right) addition of an equal volume of 200 mM sucrose in imaging buffer. These recordings were performed at 37°C using a spinning disk confocal microscope (compare vs. room temperature widefield imaging, Fig 3). N=14 FOVs (0 mM), 21 FOVs (100 mM). Horizontal lines denote the mean for each condition. *:p<0.05, unpaired 2-sample t-test.

**Supplemental Figure 7.**
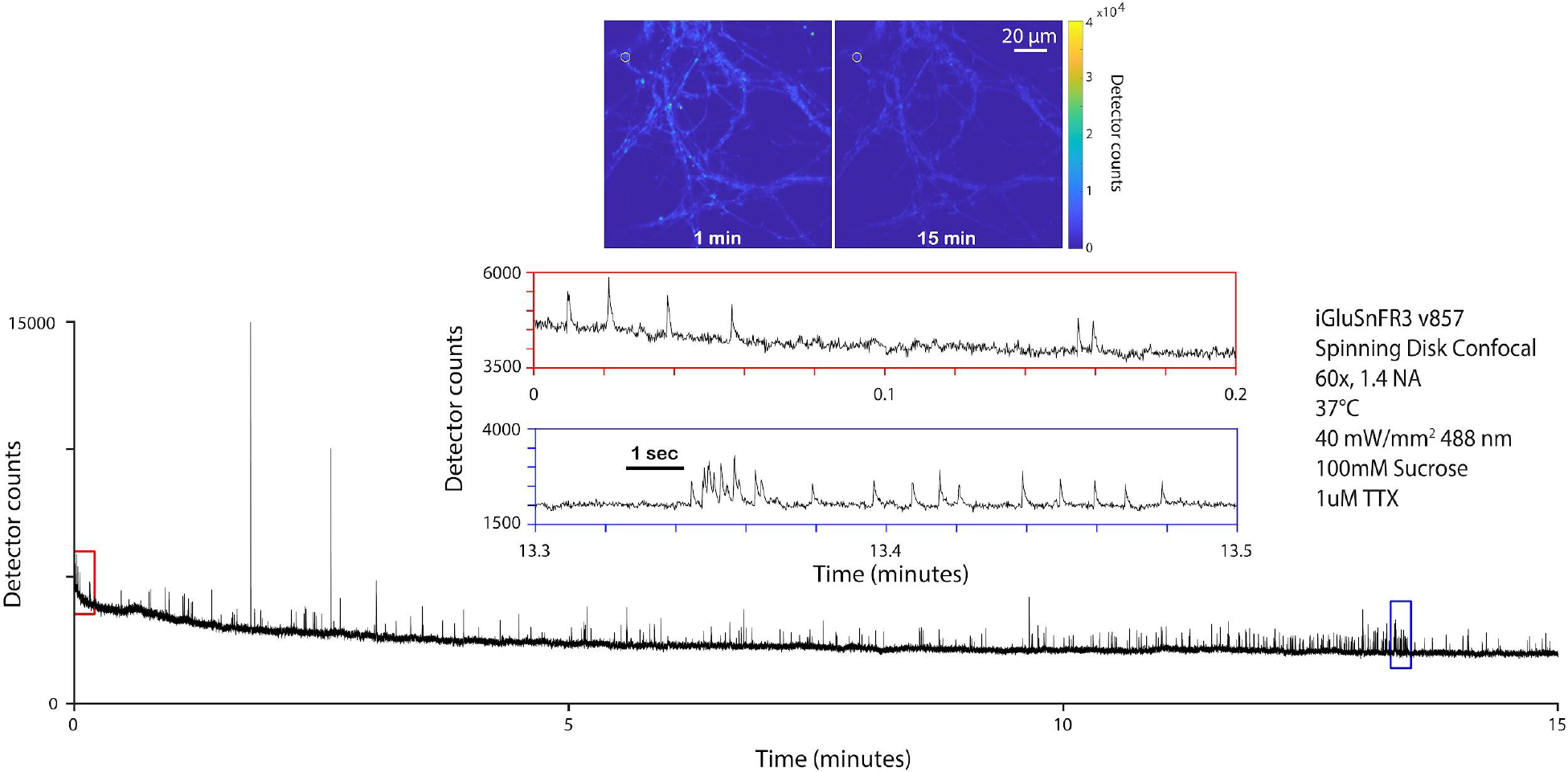
Long-duration recording of optical minis with iGluSnFR3.v857. Primary culture expressing iGluSnFR3.v857 as recorded with a spinning disk confocal microscope. (top) Inset images show frames from the beginning and end of the recording on the same intensity scale. Yellow circle denotes the site for which the traces below are plotted. Bottom, raw intensity trace for the highlighted site. Red and blue insets show zoom of highlighted regions at beginning and end of the recording, respectively. Despite moderate photobleaching over the long recording period, the SNR does not decrease over 15 minutes of recording, potentially due to bleaching of unproductive fluorescent background.

**Supplemental Figure 8.**
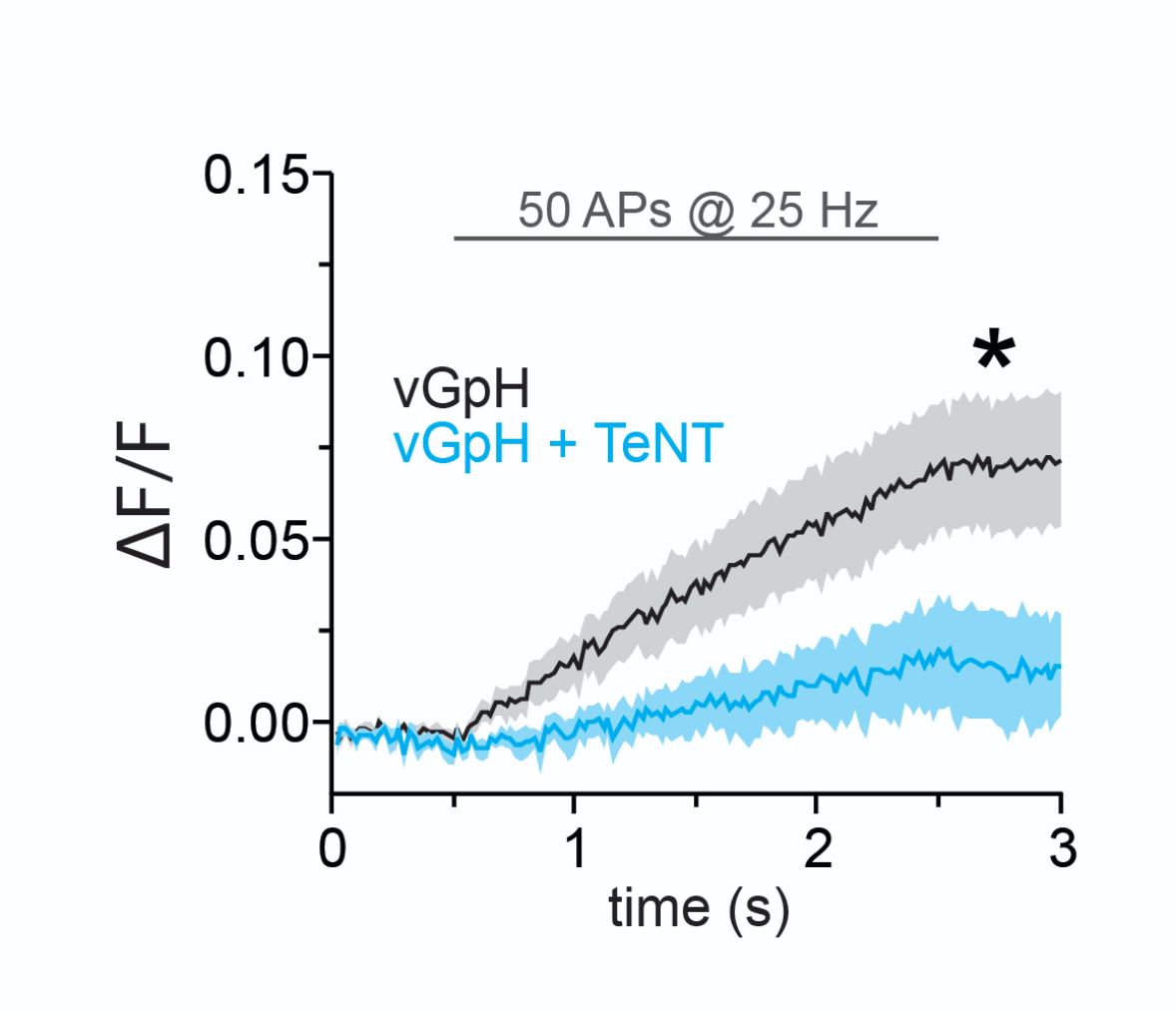
Effect of TeNT on exocytosis measured using a genetically encoded pH indicator (pHluorin) fused to a vesicular glutamate transporter (vGpH). Control (black; 7 cells) and TeNT-LC (cyan; 12 cells) expressing cells were imaged during field stimulation with 50 APs at 25 Hz. Exocytosis values are normalized to NH_4_Cl treatment as previously described (Sankaranarayanan, S., De Angelis, D., Rothman, J.E., and Ryan, T.A. (2000)). Lines denote mean, shaded areas denote SEM. * p<0.05, unpaired Student’s t-test.

**Supplemental Figure 9.**
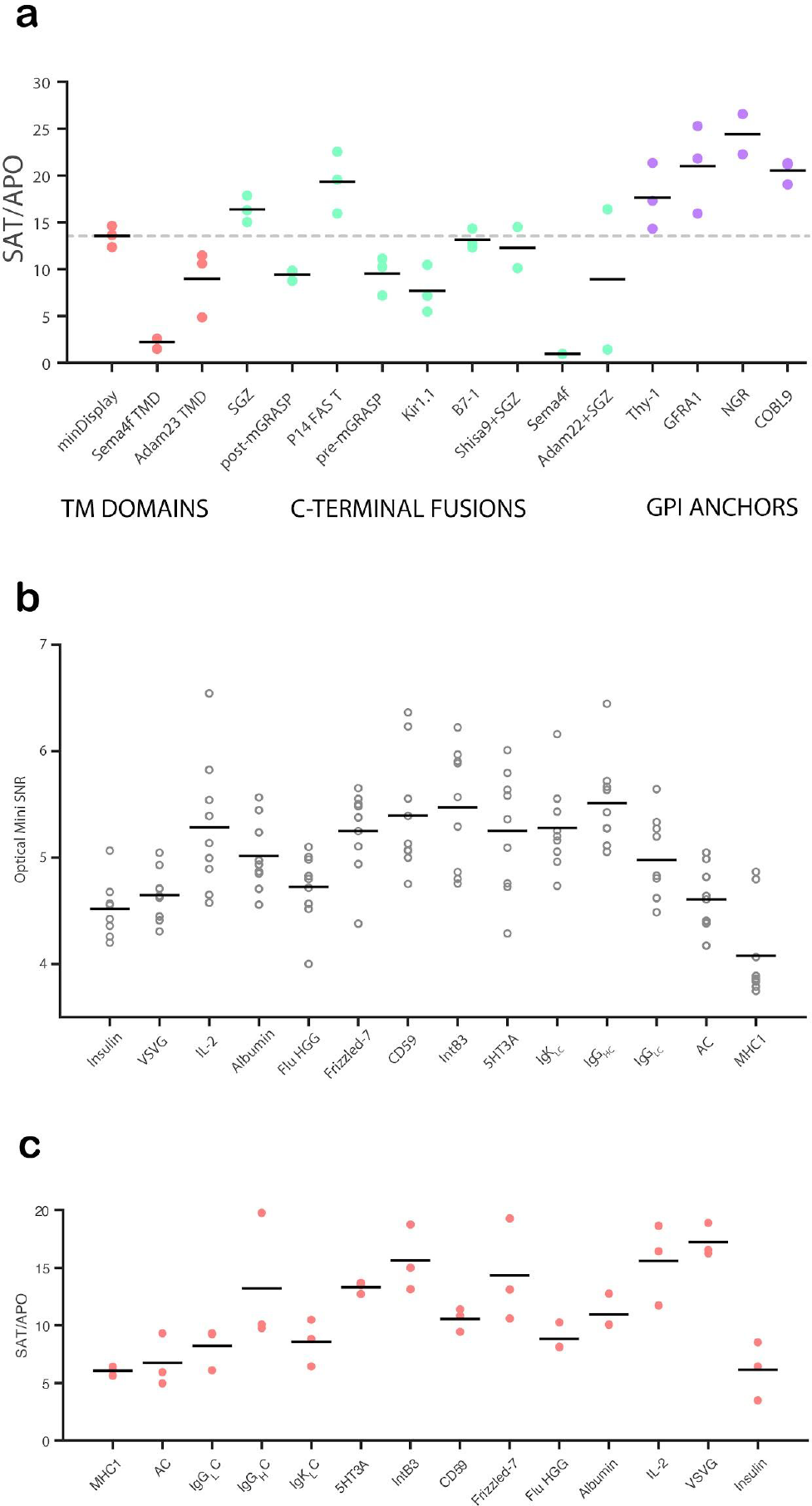
Characterization of surface display construct variants. a) Fold brightness change of iGluSnFR3.v857 in different display construct variants upon addition of saturating glutamate (20 mM). Data points denote individual culture wells. b) Mean SNR of optical minis for v857.GPI in the GPI backbone with different N-terminal leader sequences. Data points denote individual fields of view. c) Fold brightness change of v857.GPI with different N-terminal leader sequences upon addition of saturating glutamate (20 mM). In all cases, black lines denote the mean for each group. Sequences for each variant are presented in Supplemental Table 2.

**Supplemental Figure 10.**
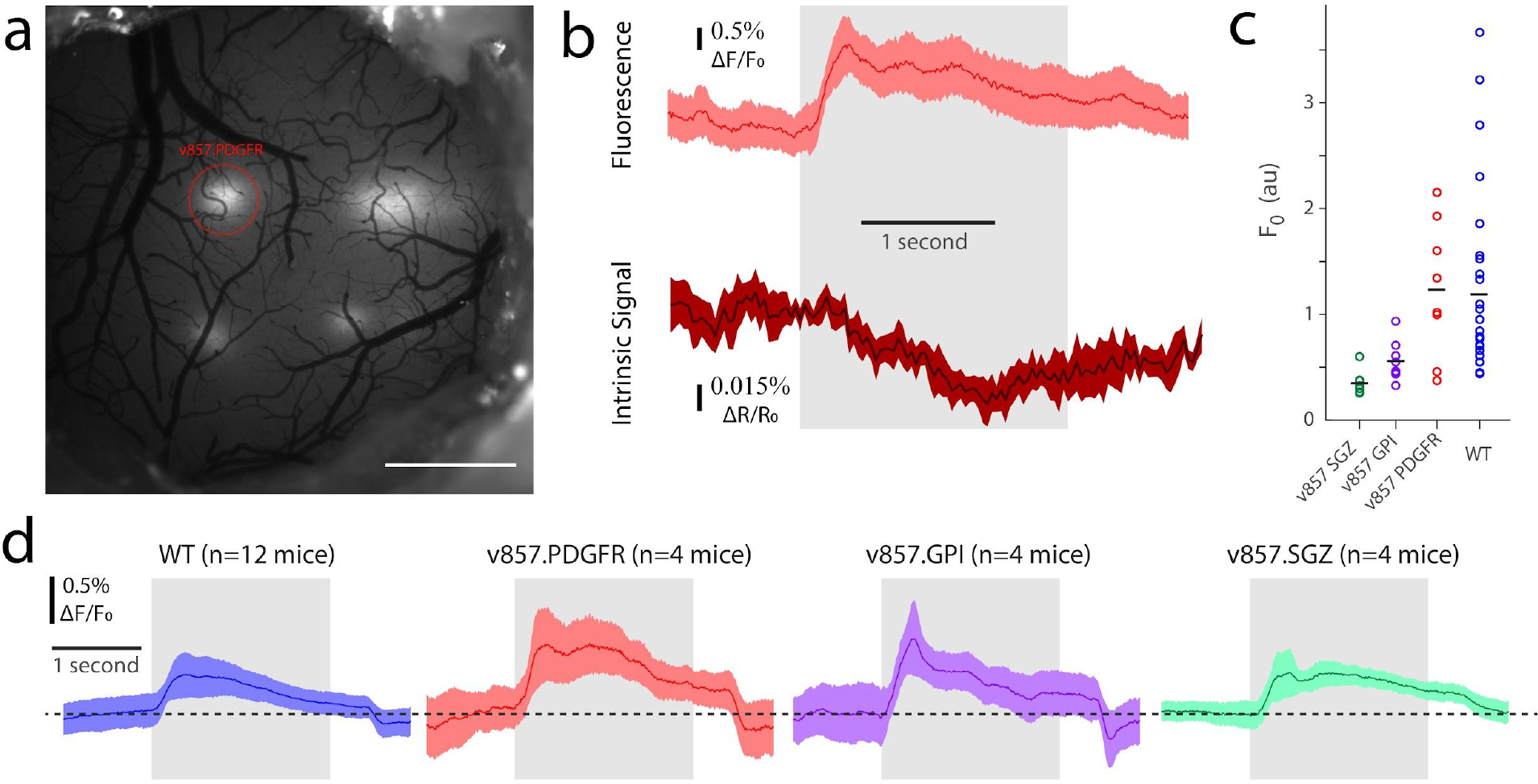
1P cortical imaging of iGluSnFR3 variants. a) Low-magnification image of mouse dorsal cortex with four injection sites. Red circle marks a region injected with v857.PDGFR. Scale bar: 1 mm. b) Mean response (shading denotes SEM) of the region highlighted in (a) to visual motion stimuli over a single session (N=64 stimuli), in fluorescence (top) and intrinsic signal (bottom). Gray region denotes time of visual stimulus. c) Baseline mean brightness of injected regions for 4 iGluSnFR variants, 3 weeks after AAV injection at matched titers. Each marker denotes a field of view, black lines denote means. d) Mean fluorescence (shading denotes SEM) response across the 50% most responsive sessions across all injection sites, for the 4 constructs. Gray region denotes time of visual stimulus.

**Supplemental Figure 11.**
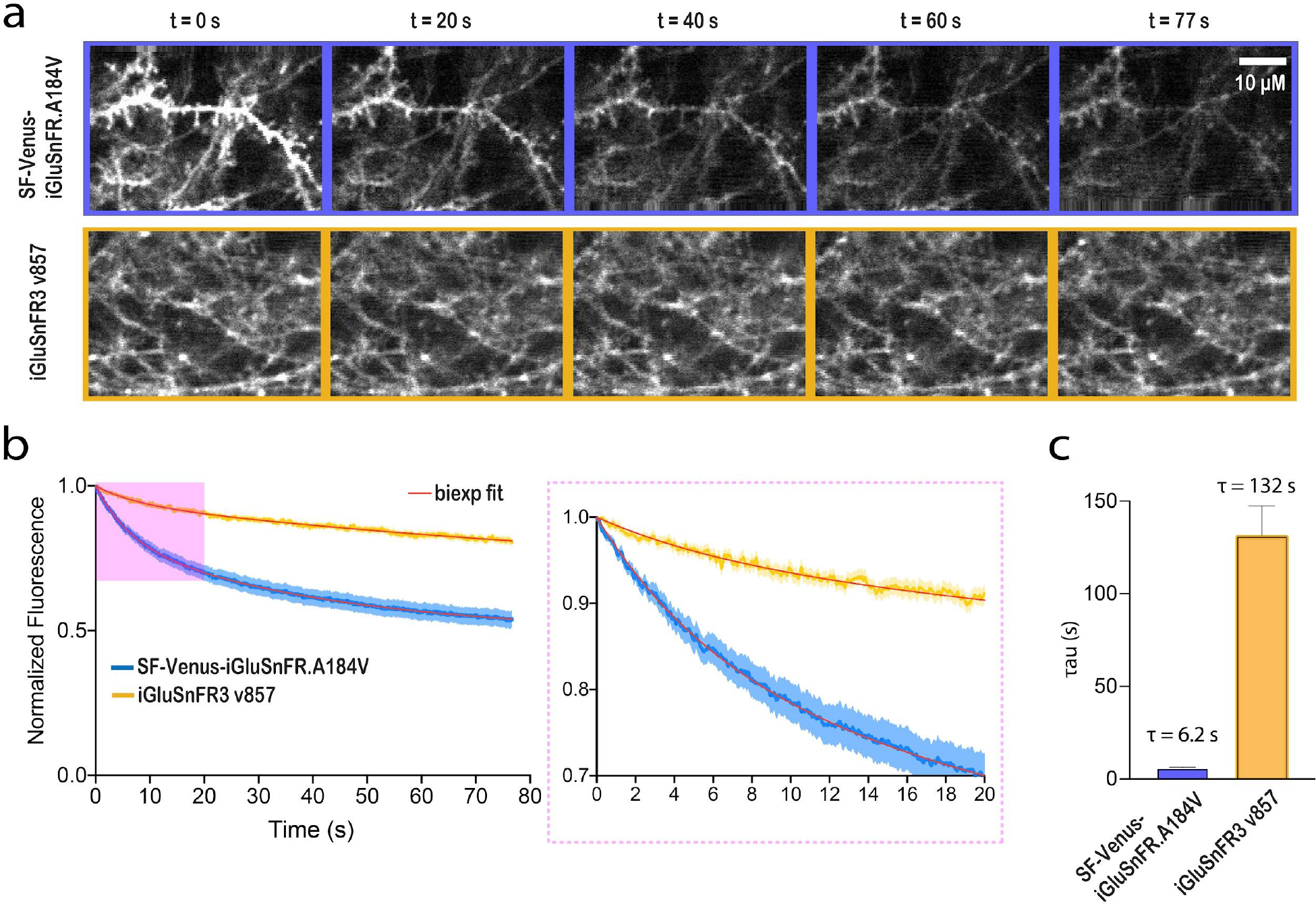
Photostability of iGluSnFR3 and SF.Venus-iGluSnFR.A184V under 2P excitation in vivo. a) Representative images of SF-Venus-iGluSnFR.A184V (blue; top) and iGluSnFR3.v857 (yellow; bottom), both in the PDGFR backbone, expressed in excitatory cortical neurons in mouse visual cortex, over the course of two-photon recordings at matched excitation power. Images are shown with the same brightness scale. b) Brightness, normalized to its initial value, of the two variants. Shading denotes SEM. A184V, n=34 FOV’s from 5 mice; v857, n=27 FOV’s from 5 mice. Curves are fit with two-phase exponential decay (red line). Right panel is a zoom-in view from 0 sec to 20 sec. c) Time constants of the fast component of the two-phase decay.

**Supplemental Figure 12.**
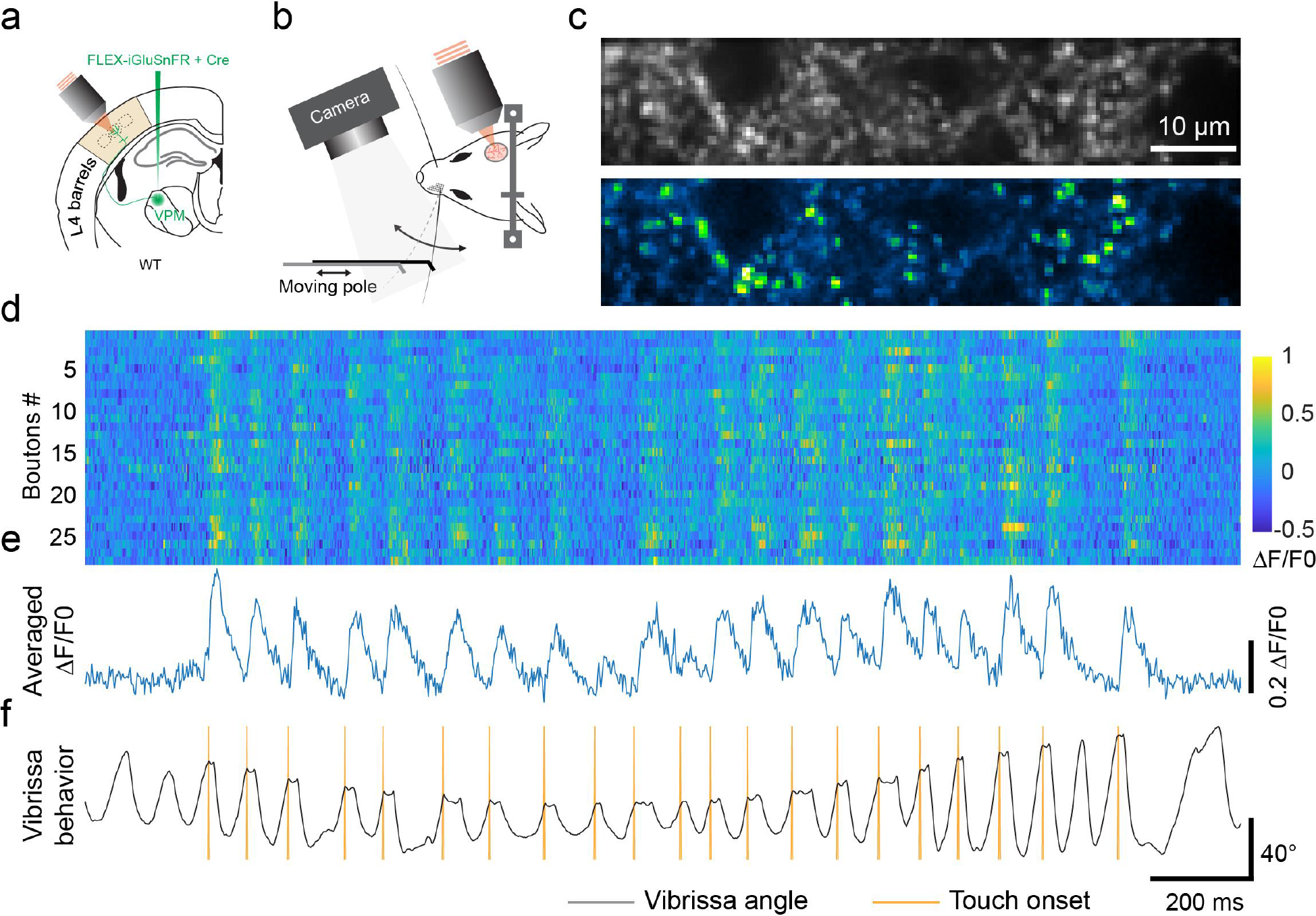
Functional in vivo imaging of layer 4 TC boutons during a pole touching task with iGluSnFR3.v857.GPI. a) Schematic of viral injection of AAV2/1.hSyn.iGluSnFR3.v857.GPI in VPM to label the TC boutons in layer 4. b) Schematic of the dynamic pole touch tasking during active sensing. The pole moves back and forth at a range of 5 mm along the azimuthal direction with an average speed of 1.25 mm/s. Images are acquired at the frame rate of ∼500 Hz. c) Average (top) and SD (bottom) of the image of TC axons and boutons labeled by v857.GPI in layer 4 of vibrissa cortex. d) Raster plot of the touch response from individual boutons detected from panel c. e) Averaged touch response across all boutons in panel d. f) Corresponding whisking trace and touch onset. This experiment was performed with a single subject and is a representative session.

## Supplemental Tables

**Supplemental Table 1.** Photophysical characterization of SF-Venus-iGluSnFR.A184V (wild-type), iGluSnFR3.v82, and iGluSnFR3.v857. All measurements are made in purified protein. N=3, error bars represent SEM.

| Variant Name | SF-Venus-iGluSnFR.A184V |  | iGluSnFR3.v82 |  | iGluSnFR3.v857 |  |
| --- | --- | --- | --- | --- | --- | --- |
|  | APO | SAT | APO | SAT | APO | SAT |
| 1-photon Excitation maxima $\lambda_{exc}$ (nm) | 512 | | 512 | | 502 | |
| 1-photon Emission maxima $\lambda_{exc}$ (nm) | 530 | | 530 | | 522 | |
| 1-photon $\Delta F/F_0$ | $2.9 \pm 0.1$ | | $13.1 \pm 0.9$ | | $54.0 \pm 2.6$ | |
| $K_d$ ($\mu M$) | $24.4 \pm 2.4$ | | $31.7 \pm 3.9$ | | $195.9 \pm 14.8$ | |
| Apparent Hill coefficient ($n_H$) | $0.97 \pm 0.096$ | | $1.1 \pm 0.1$ | | $1.2 \pm 0.097$ | |
| Apparent pKa | $7.48 \pm 0.08$ | $6.63 \pm 0.06$ | $7.88 \pm 0.12$ | $6.61 \pm 0.06$ | $6.88 \pm 0.26$ | $6.74 \pm 0.08$ |
| $K_M$ ($\mu M$) | $180 \pm 69$ | | $1960 \pm 210$ | | $16800 \pm 1550$ | |
| Extinction Coefficient $\epsilon$ ($M^{-1}cm^{-1}$) | $34,000 \pm 990$ | $116,000 \pm 3300$ | $11,800 \pm 840$ | $105,000 \pm 1600$ | $2,000 \pm 540$ | $40,000 \pm 2900$ |
| Quantum Yield $\Phi$ | $0.54 \pm 0.03$ | $0.54 \pm 0.01$ | $0.41 \pm 0.02$ | $0.60 \pm 0.01$ | $0.26 \pm 0.03$ | $0.93 \pm 0.03$ |
| 1-photon Brightness (x1000) | 18.4 | 62.6 | 4.8 | 63.0 | 0.5 | 37.2 |
| 2-photon $\Delta F/F_0$, (950 nm / 1030 nm) | 2.67 / 2.97 | | 9.81 / 10.44 | | 61.54 / 67.08 | |
| 2-photon Brightness $F_2$ (GM), (950 nm / 1030 nm) | 25.6 / 25.0 | | 21.1 / 19.9 | | 22.6 / 8.16 | |

**Supplemental Table 2.**
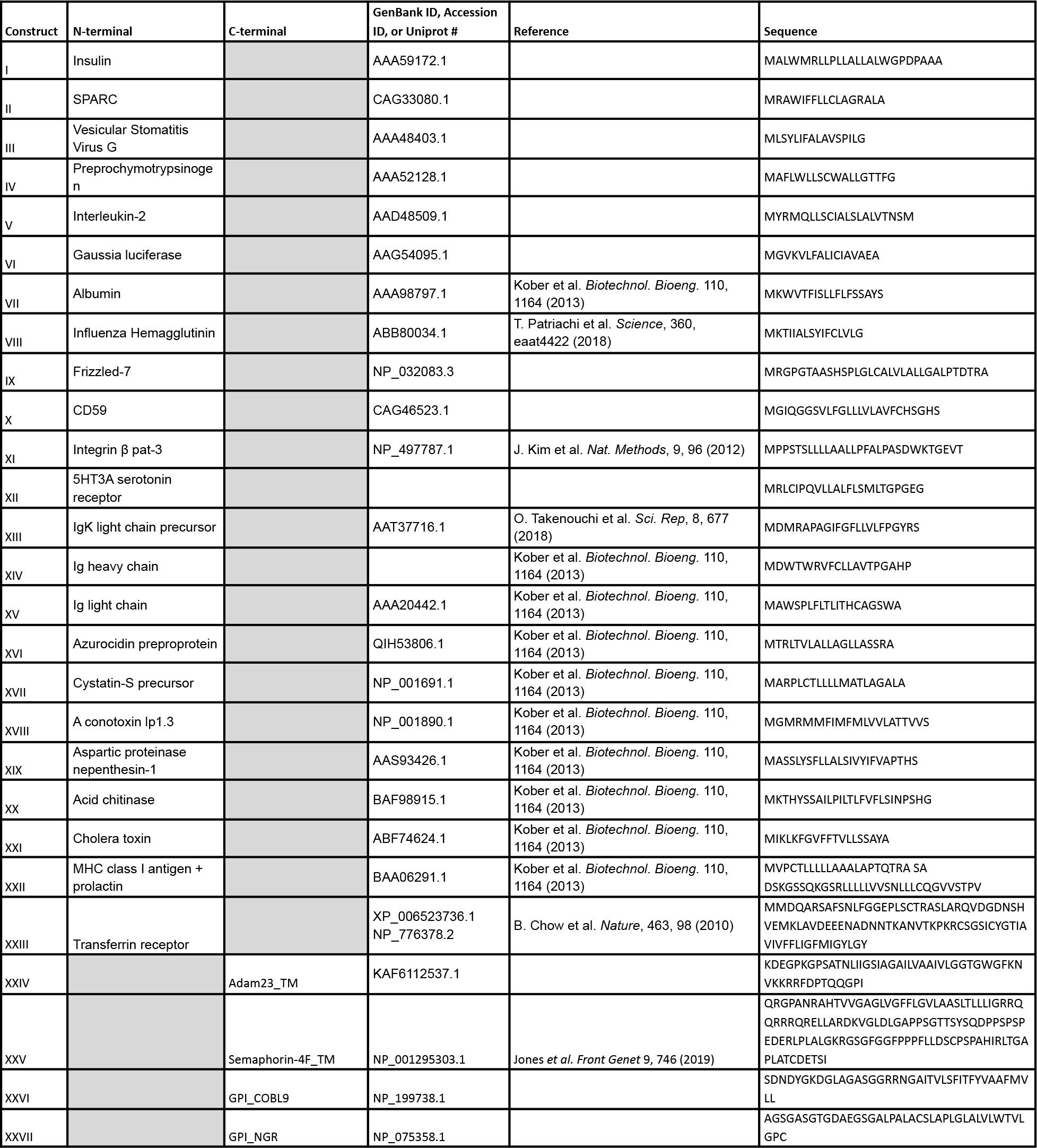
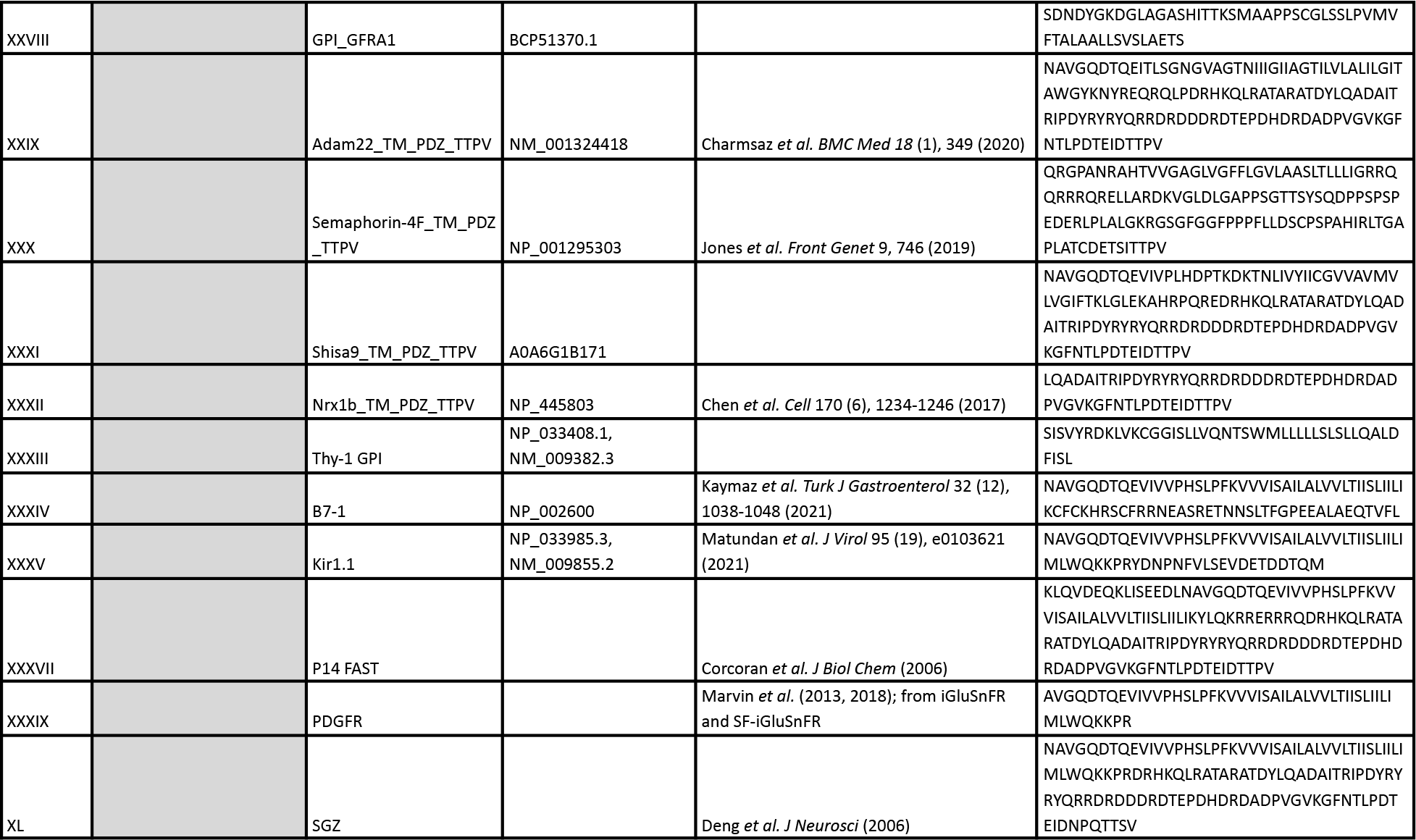
Sequences of surface display construct variants used in the N-terminal screen and C-terminal screen. Sequences I to XXIII show the protein sequences that were used for the N-terminal screen (see Supplemental Figure 9), and constructs XXIV to XL show the amino acid sequences for the C-terminal screen (see Figure 4). N-terminal mutations were introduced relative to iGluSnFR3.v857.GPI (sequence XXVI). C-terminal mutations were introduced relative to iGluSnFR.v857 with the IgK N-terminal secretion leader (sequence XIII). All sequences were codon-optimized for *Mus musculus*.

**Supplemental Table 3.** Summary of iGluSnFR variants tested in this study. ‘WT’ refers to SF-Venus-iGluSnFR.A184V (Marvin *et al* 2018).

| Variants tested in this study | Reference | Figures | Supplemental Figures |
| --- | --- | --- | --- |
| SF-iGluSnFR.A184V | *Marvin et al (2018)* | 3, 6 |  |
| SF-iGluSnFR.A184S | *Marvin et al (2018)* | 5 |  |
| SF-Venus-iGluSnFR.A184V | *Marvin et al (2018)* | 1, 2, 6 | 1, 2, 3, 4, 5, 10, 11 |
| SF-Venus-iGluSnFR.A184S | *Marvin et al (2018)* | 1 |  |
| iGluSnFR3.v82 | *This study* | 1, 2, 6 | 1, 2, 3, 4, 5 |
| iGluSnFR3.v857 | *This study* | 1, 2, 3, 4, 5, 6 | 1, 2, 3, 4, 5, 6, 7, 9, 10, 11, 12 |

## Supplemental Videos

**Supplemental Video 1.** Widefield recording of optical minis using iGluSnFR3.v857.GPI in mixed hippocampal and neocortical primary cultures 14 days post transfection. Culture medium was exchanged with hyperosmotic buffer containing 100 mM sucrose, and action potentials were silenced using 2 µM TTX. Video is recorded at 100 frames/second using a Nikon Eclipse microscope (60X 0.4NA).

**Supplemental Video 2.** *In vivo* 2-photon recording of spontaneous activity (no visual stimulus) in excitatory cortical neurons in mouse visual cortex using iGluSnFR3.v857.PDGFR using a custom 2-photon microscope (20X, 1.0NA, Exc - 1010 nm, Em 525/50, 156 Hz).

**Supplemental Video 3.** Averaged images of thalamocortical boutons with sparse labeling of iGluSnFR3.v857.PDGFR in L4 during whisker stimulation. Images are acquired at the frame rate of 247.9 Hz for 60 s and interpolated to 500 Hz and averaged to 1s. Images are acquired at the depth of 350 µm below pia with a field of view of 50x25 µm.

**Supplemental Video 4.** Averaged images of dendritic spines labeled with iGluSnFR3.v857.GPI in L4 during whisker stimulation. Images are acquired at the frame rate of 130 Hz for 60 s and interpolated to 500 Hz and averaged to 1s. Images are acquired at the depth of 365 µm below pia with a field of view of 50x30 µm.

## Footnotes

1 This numbering scheme uses position indices 2 lower than those used by Marvin et al 2013

